# Chronic evoked seizures in young pre-symptomatic APP/PS1 mice induce serotonin changes and accelerate onset of Alzheimer’s disease-related neuropathology

**DOI:** 10.1101/2023.01.05.522897

**Authors:** Aaron del Pozo, Kevin M. Knox, Leanne M. Lehmann, Stephanie Davidson, Seongheon Leo Rho, Suman Jayadev, Melissa Barker-Haliski

## Abstract

**Objective:** People with early-onset Alzheimer’s disease (AD) are at elevated seizure risk. Further, chronic seizures in pre-symptomatic stages may disrupt serotonin pathway-related protein expression, precipitating the onset of AD-related pathology and burden of neuropsychiatric comorbidities.

**Methods:** 2-3-month-old APP/PS1, PSEN2-N141I, and transgenic control mice were sham or corneal kindled for 2 weeks to model chronic seizures. Seizure-induced changes in glia, serotonin pathway proteins, and amyloid β levels in hippocampus and prefrontal cortex were quantified.

**Results:** APP/PS1 mice experienced worsened mortality versus kindled Tg-controls. APP/PS1 females were also more susceptible to chronic kindled seizures. These changes correlated with a marked downregulation of hippocampal tryptophan hydroxylase 2 and monoamine oxidase A protein expression compared to controls; these changes were not detected in PSEN2-N141I mice. Kindled APP/PS1 mice exhibited amyloid β overexpression and glial overactivity without plaque deposition. PSEN2 protein expression was AD model-dependent.

**Significance:** Seizures evoked in pre-symptomatic APP/PS1 mice promotes premature mortality in the absence of pathological Aβ deposition. Disruptions in serotonin pathway metabolism are associated with increased glial reactivity and PSEN2 downregulation without amyloid β deposition. This study provides the first direct evidence that seizures occurring prior to amyloid β plaque accumulation worsen disease burden in an AD genotype-specific manner.

**Highlights:**

- Seizures are a comorbidity in Alzheimer’s disease that may worsen disease burden.
- Pathological overlap between both neurological disorders is understudied.
- Young APP/PS1, but not PSEN2-N141I mice, have increased seizure-induced mortality.
- Seizures reduce hippocampal serotonin pathway proteins only in young APP/PS1 mice.
- Kindled young APP/PS1 mice have glial hyperactivity before amyloid β accumulation.

## INTRODUCTION

Alzheimer’s disease (AD) is the most common cause of dementia affecting more than 55 million people worldwide and is characterized by pathological amyloid β (Aβ) accumulation (Tort-Merino et al., 2022; Yang et al., 2022). Despite being an aging-related disorder, 5% of cases occur prior to age 65 and are thus considered early-onset AD (EOAD) (Tort-Merino et al., 2022; Yang et al., 2022). EOAD can arise due to duplications or genetic variants in three key deterministic risk genes – amyloid precursor protein (*APP*), and the homologous presenilin 1 (*PSEN1*) and 2 (*PSEN2*) genes. EOAD shares the same essential neuropathological traits as late-onset AD (i.e., Aβ plaques and neurofibrillary tangles) but differs in several features (Tort-Merino et al., 2022; Yang et al., 2022). For example, EOAD may have a more aggressive course at both clinical and neuropathological levels, with myoclonic and focal seizures emerging in recent years as an understudied comorbidity in AD (Lehmann et al., 2021; Tort-Merino et al., 2022; Yang et al., 2022). Notably, untreated seizures may accelerate AD-associated cognitive decline and neuropathological hallmarks (Friedman et al., 2012). The relative risk of unprovoked seizures markedly increases in AD, reaching up to 87-fold greater risk for seizures in individuals with EOAD onset between 50 and 59 years versus that of the general population (Lehmann et al., 2021). Thus, untreated chronic seizures likely worsen AD trajectory.

Epilepsy and AD share many pathological similarities: temporal lobe atrophy, neuronal death, gliosis, neuritic alterations, and neuroinflammation (Lehmann et al., 2021). Particularly, diverse studies demonstrated that both AD (Abbink et al., 2020) and epilepsy (Sano et al., 2021) subjects exhibited an increase in astrocyte and microglial reactivity. More critically, seizures in AD potentially reduce quantity of life. Mortality specifically from AD in patients with epilepsy has increased over 200% in the last 20 years, whereas patients with epilepsy alone have seen a significant (27%) decrease in mortality (DeGiorgio et al., 2020; Zawar & Kapur, 2023). Thus, untreated seizures in AD represent a pressing public health concern to mitigate the functional disease burden and potential excess mortality associated with AD.

While the mechanisms connecting the presence of seizures and associated mortality risk in AD remain unknown, serotonin (5-HT) is hypothesized to play a key role in regulating epilepsy-associated premature mortality. Preclinical sudden unexpected death in epilepsy (SUDEP) models suggests that a genetic or acquired defect in the 5-HT system increases seizure-induced mortality risk (Buchanan et al., 2014; Massey et al., 2021). 5-HT neurons project to all the major respiratory nuclei, including the dorsal raphe (Ptak et al., 2009; Turkin et al., 2021). They regulate many brain functions critical to post-ictal respiratory control, such as stimulating respiratory output (Buchanan et al., 2014), acting as chemoreceptors that respond to an increase in systemic CO_2_ levels (Buchanan et al., 2014), and enabling plasticity of the respiratory network in response to intermittent hypoxia (Buchanan et al., 2014; Li & Buchanan, 2019; Petrucci et al., 2020). Lesions of 5-HT neurons can decrease respiratory output. Thus, chronic seizure-evoked 5-HT dysregulation may increase propensity for premature mortality due to decreased autoresuscitation capacity and respiratory drive (Buchanan et al., 2014; Li & Buchanan, 2019; Petrucci et al., 2020).

Postmortem AD studies also show reduced CNS 5-HT levels (Lin et al., 2015; Metaxas et al., 2019; Tejani-Butt et al., 1995; von Linstow et al., 2022), including in the dorsal raphe and hippocampus (HPC; (Tejani-Butt et al., 1995)). Advanced age (18-24-month-old) APP/PS1 mice have decreased serotonin transporter (SERT) density in the parietal and frontal cortex and Aβ40 itself can further reduce SERT activity (Metaxas et al., 2019). Additionally, genetic deletion of tryptophan hydroxylase 2 (TPH2) expression in the AD-associated APP/PS1 mouse model disproportionally increases mortality, suggesting an understudied interaction between AD-related genotypes and dysregulation of 5-HT synthesis that exacerbates the risk for premature mortality (von Linstow et al., 2022). These prior clinical and preclinical studies largely focused on older individuals at symptomatic stages, i.e., the age when AD-related cognitive symptoms are present and dense core Aβ plaques are evident, making it difficult to identify novel contributing mechanisms. No study has yet directly assessed the seizure-induced changes in 5-HT system regulation prior to Aβ plaques deposition to further uncover the bidirectional interactions between seizures and AD risk factors on susceptibility to premature mortality.

While APP/PS1 models are most commonly used to assess the functional and pathological impacts of evoked and spontaneous seizures in AD (Vande Vyver et al., 2022b; Ziyatdinova et al., 2011; Ziyatdinova et al., 2015), PSEN2 variant models are also useful to assess the biological heterogeneity of AD pathology (Fung et al., 2020; Jayadev et al., 2013; Jayadev, Leverenz, et al., 2010), including seizure risk (Beckman et al., 2020) and seizure-evoked cognitive deficits (Knox et al 2023). Individuals with the most common EOAD PSEN2 variant, N141I, experience seizures and this variant may increase inflammatory response (Fung et al., 2020; Jayadev et al., 2013) that could itself influence seizure susceptibility (Vezzani, 2005; Vezzani et al., 2012). PSEN2-N141I models are also relevant to study the non-neuronal contributions to AD without Aβ plaque accumulation (Fung et al., 2020). We have previously demonstrated that young APP/PS1 mice establish a hyperexcitable neuronal network faster than non-transgenic mice and that chronic seizures increase premature mortality (Vande Vyver et al., 2022b). We thus quantified the seizure susceptibility and seizure-induced 5-HT system changes in young-adult (2-3 months-old) APP/PS1 and PSEN2-N141I EOAD mouse models to define the extent to which chronic seizures evoked prior to potential neuropathological alterations conferred genotype-specific effects on seizure susceptibility, mortality, 5-HT system regulation, and neuroinflammation. While APP/PS1 mice are known to demonstrate modest increases in premature mortality in later life (Minkeviciene et al., 2009), we hypothesized that evoked seizures and accompanying accelerated mortality in pre-symptomatic stages reflected potential dysfunction in 5-HT system regulation in a manner consistent with that which is observed in genetic epilepsy models with a SUDEP phenotype. This study critically reveals that uncontrolled seizures occurring prior to pathological Aβ accumulation can accelerate the functional and pathological burden of AD in a genotype-specific manner.

## Material and Methods

### Animals

Male APP^swe^/PS1^dE9^ mice (strain 034832-JAX) were kindly provided by Dr. Gwenn Garden from stock originally purchased from the Jackson Laboratory (Bar Harbor, ME). PCR-confirmed male APP/PS1 mice were bred to wild-type female C57Bl/6J mice. PSEN2-N141I transgenic mice and their non-excised transgene controls were kindly provided by Dr. Suman Jayadev, with breeding as previously described (Fung et al., 2020).

Mice were housed on a 14:10 light cycle (on at 6 h00; off at 20 h00) in ventilated cages with free access to irradiated food and water, as previously described (Meeker et al., 2019). Housing conditions conformed to the *Guide for the Care and Use of Laboratory Animals* and all animal work was approved by the UW Institutional Animal Care and Use Committee (protocol 4387-01) and conformed to ARRIVE guidelines. All behavioral testing was performed between 9 h00 and 17 h00 by an experimenter blinded to genotype.

### Corneal kindling

Young adult male and female (2-months-old at kindling initiation) APP/PS1 and PSEN2-N141I versus their respective transgene negative (Tg-) control mice were corneal kindled with a 3 sec, 60 Hz 1.6-2.0 mA bilateral sinusoidal pulse twice per day for 2-3 weeks, consistent with our prior reports (Barker-Haliski et al., 2016; Beckman et al., 2020; Knox KM, 2023). Sham kindled mice of each genotype were similarly handled at each twice daily stimulation session, but no electrical current was delivered (Supplemental Figure 1A). Stimulation intensity was confirmed in a separate cohort of mice and based on the subconvulsive minimal clonic seizure thresholds for age- and sex-matched AD-associated mice (Beckman et al., 2020).

Kindled seizure severity was scored by an investigator blinded to genotype according to the Racine scale (Racine et al., 1972). Group sizes and the total number of mice for each group to achieve kindling criterion of 5 consecutive Racine stage 5 seizures are detailed in Supplemental Table 1.

### Mortality assessment

Percentage of mice/group and kindling sessions elapsed before death was used to quantify the excess mortality of both AD phenotypes subjected to corneal kindling. Animals that prematurely died during kindling were post-hoc sorted into two categories: death before or after kindling acquisition. Mice classified as “before” were those that died prior to achieving fully kindled criterion (five consecutive stage 5 seizures). Mice that were euthanized for terminal collections on the study end date were considered to have met study endpoint.

### Euthanasia and terminal sample collections

Mice were kindled for 10-14 days and then terminal samples collected by live decapitation 24-72 h after attaining kindling criterion. Mice were euthanized by live decapitation 24-72 hours after reaching corneal kindling criterion. Brain was rapidly excised and hemisectioned along the sagittal plane. The right HPC and prefrontal cortex (PFC) were microdissected and flash frozen to evaluate the protein changes secondary to AD genotype and/or corneal kindling via western blot (WB). The left hemisphere was flash frozen in 2-methylbutane on dry ice 24h post PFA immersion for use in immunohistochemistry (IHC) analysis of posterior parietal association cortex (region overlying CA1 (CTX)), CA1, CA3, and dentate gyrus (DG) of dorsal HPC.

### Preparation of biological samples

A subset of mice were used to measure the molecular changes induced by corneal kindling. Sexes were pooled for biomolecular studies in kindled and sham kindled animals within each genotype because the primary outcome of kindled seizure status (or sham) was the same between sexes.

### Western blot

For WB, HPC and PFC samples were homogenized in tissue protein extraction reagent (TPER) [10 mL/g] and protease inhibitor cocktail (Millipore Sigma) [10 µL/mL]. The homogenization frequency was 50 oscillations for 4 min, followed by centrifugation at 12 000× g for 10 min at 4°C. Total protein levels of the area of interest were measured using the BCA method based on the principle of protein-dye binding. After adjusting protein levels following the BCA protocol, homogenates were mixed with Laemmli sample buffer (Bio-Rad®) containing β-mercaptoethanol (50 μL/mL of Laemmli), to get a final concentration of 1 mg/mL, and 20 uL was loaded into an electrophoresis gel. Proteins were blotted onto a nitrocellulose membrane (Bio-Rad®) with a dry transfer system (Bio-Rad), incubated with specific primary antibodies (Supplemental Table 2). After that, each membrane was incubated with their respective secondary antibody and protein bands of interest revealed by use of the AP-conjugate colorimetric protocol. Blots were digitized and band intensities quantified by densitometric analysis using ImageJ (NIH ImageJ® software, National Biosciences, Lincoln, NE, USA). Raw densitometric data in different blots were transformed as fold change of the control mean, expressed in arbitrary units of OD, and Ponceau red was used as loading controls (Romero-Calvo et al., 2010).

### Immunohistochemistry

Left hemispheres were immersed in PFA 4% for 24 h. Then, brains were submerged in 30% sucrose. 24 h later, brains were flash frozen and store at -80°C. Two consecutive 20 µm-thick sections/mouse from the dorsal HPC (anteroposterior [AP] from Bregma: −1.34 mm) and ventral HPC (AP from Bregma: −2.24 mm) were sectioned on a cryostat (Leica DM1860) and slide-mounted. Tissues were processed for semi-quantitative immunohistochemistry for astrocytes (glial fibrillary acidic protein; GFAP), microglia (Iba-1), neurons (NeuN) and β-amyloid (6e10 and Thioflavin S) (Supplemental Table 2). Briefly, GFAP directly conjugated to Cy3 fluorophore (1:1000; C9205 Sigma-Aldrich), Iba1, NeuN, Thioflavin and 6e10 antibodies were applied under 200 μL coverwells in a 5% goat serum in 1 × phosphate-buffered saline (PBS) solution overnight at 4°C according to previously published methods (Knox et al., 2021; Loewen et al., 2016). Brains were incubated for 2 h in their respective secondary antibody. Finally, all slides were counterstained with 4’,6-diamidino-2-phenylindole (DAPI) (D9542, 1100, Sigma Aldrich). Slides were then rinsed and coverslipped with Prolong Gold antifade reagent. Slides were imaged on an upright fluorescent microscope (Leica DM-4) with a 5x objective (20x final magnification) with constant acquisition settings. Photomicrographs (n = 4 brain sections/mouse) were analyzed with Leica Thunder software and visually scored by a blinded investigator for the presence of positive cells in CTX, CA1, CA3, and DG of dorsal HPC. For quantification, we measured the mean density of immunolabelling in the selected areas as previously published.

### Statistical analysis

The sample sizes in the different experimental groups were always ≥5 (Supplemental Table 1). Mean seizure score during the corneal kindling period was assessed within APP/PS1 and PSEN2-N141I transgenic mice and matched Tg controls using Friedman’s test. Latency to attain corneal kindling criterion and survival rate during kindling were assessed using a Mantel-Cox Log-Rank test and presented as a Kaplan-Meier curve. Seizure burden (the average of the seizure score until the end of the study) and the number of sessions to achieve the kindled criterion were assessed by a T-test and Welch correction. Molecular analyses from WB and IHC studies were assessed using one-way or two-way ANOVA followed by the Student-Newman-Keuls test or the Bonferroni test, as appropriate, using GraphPad Prism, v8.0 or later (GraphPad Software, San Diego, CA, USA). Statistical significance was defined as p < 0.05 for all tests.

## Results

### Female APP/PS1 mice are highly susceptible to 60 Hz kindled seizures

A primary goal of this study was to compare the 60 Hz kindled seizure susceptibility in two AD mouse models prior to the age when Aβ accumulation is widespread, thereby extending our earlier published works with PSEN2 null (Beckman et al., 2020; Knox KM, 2023) and APP/PS1 mice (Vande Vyver et al., 2022a). This effort was largely aimed at addressing how AD-related genotypes differentially influence seizure risk in young animals. Consistent with our 6 Hz corneal kindling studies (Vande Vyver et al., 2022a), 2-month-old APP/PS1 females kindled significantly faster compared to their respective Tg-littermates (Figure 1A, B). Moreover, APP/PS1 females also showed a 20% increase in seizure burden (Figure 1C) and needed significantly fewer sessions to achieve kindling criterion (Figure 1D). APP/PS1 males did not show accelerated kindling acquisition versus their respective Tg-littermates (Figure 1E, F). In accordance with this result, APP/PS1 males did not experience any differences in seizure burden (Figure 1G) and did not have differences in the number of sessions to achieve criterion (Figure 1H). Thus, there were marked sex-related differences in corneal kindling rates in APP/PS1 mice.

**Figure 1:**
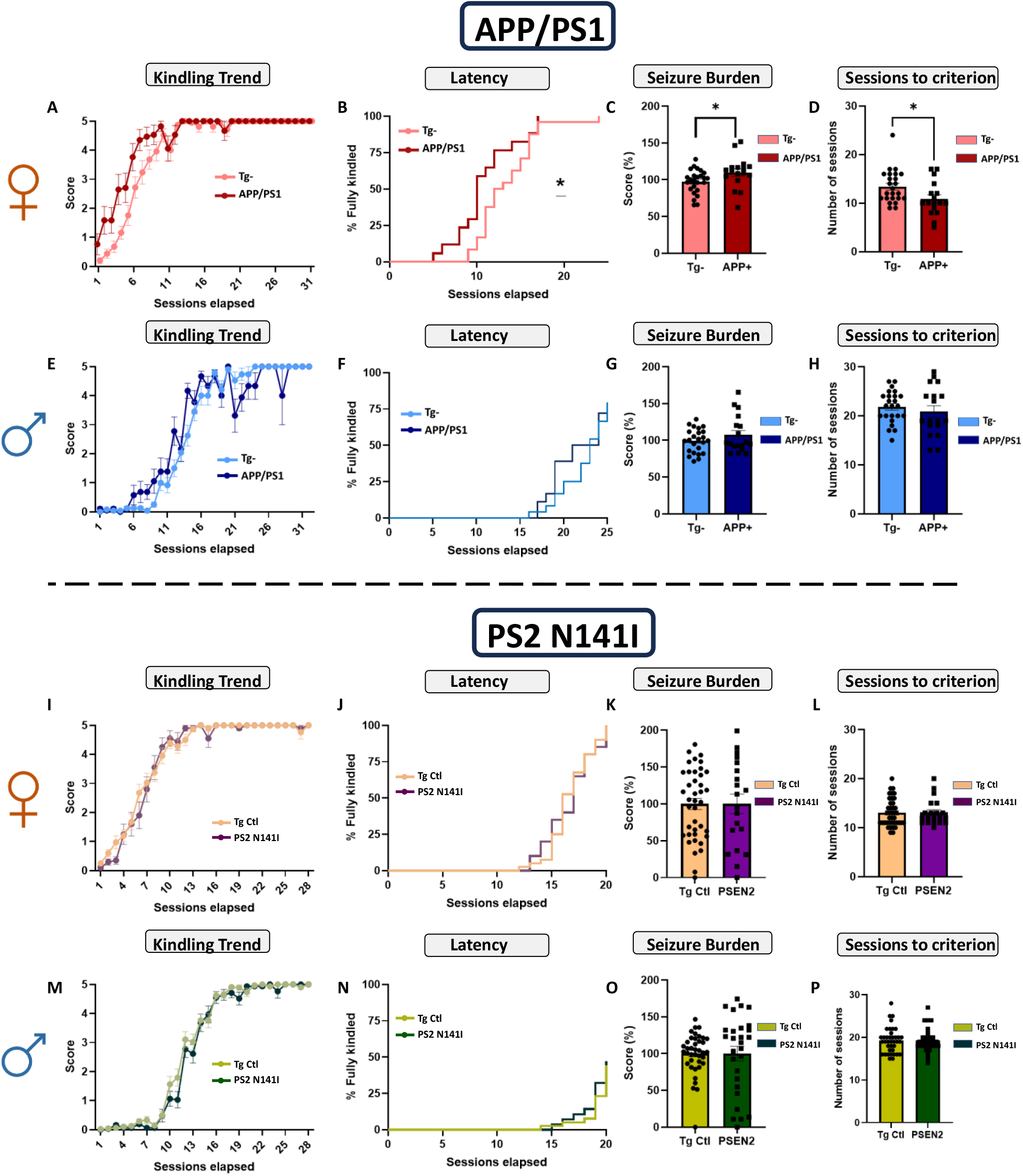
AD-associated genotype differentially influences seizure susceptibility and 60 Hz corneal kindling acquisition rate of 2-3-month-old male and female mice. APP/PS1 female mice subjected to 60 Hz corneal kindling attained A-B) criterion significantly faster than matched Tg-littermate females, C) higher seizure burden, and D) required fewer sessions to achieve kindling criterion (five consecutive Racine stage 5 seizures). Male APP/PS1 subjected to 60 Hz corneal kindling did not attain E-F) kindling criterion differences compared with matched Tg-littermate males. They did not demonstrate significant differences in G) seizure burden, or H) sessions to achieve kindled criterion. Conversely, PSEN2-N141I variant mice in both sexes did not significantly differ from transgenic control mice across all outcome measures, reflecting substantial differences in seizure susceptibility with AD-associated genotypes (I-P). (A, E, I, M) Corneal kindling acquisition rates are presented as mean seizure score ± SEM. Friedman’s test was performed to evaluate differences in mean seizure score between genotypes. (*) *p < 0.05 vs WT*. (B, F, J, N). Latency to attain kindling criterion is presented as a Kaplan-Meier plot and assessed by Log-Rank Mantel Cox test. (C, G, K, O) Seizure burden and (D, H, L, P) sessions to achieve criterion are presented as the median and the SEM and analyzed by Mann–Whitney test. (*) *p < 0.05 vs WT*. N= 15-30 mice per group.

The PSEN2-N141I model of AD is associated with a pro-inflammatory microglial response following inflammatory stimulus, such as LPS challenge (Fung et al., 2020), leading us to hypothesize that these mice would be more susceptible to kindling due to the altered microglial phenotype and reactivity on seizure risk (Eyo et al., 2017). Contrary to our hypothesis and in contrast to our earlier studies in PSEN2 null mice (Beckman et al., 2020; Knox KM, 2023), both male and female PSEN2-N141 mice did not show any significant variation in kindling acquisition rate versus their respective Tg-control mice (Figure 1I-J, M-N). There were also no significant changes in the seizure burden and stimulation sessions to achieve kindling criterion in either male or female PSEN2-N141I mice versus Tg-control mice (Figure 1K-L, O-P). Thus, kindled seizure susceptibility is highly genotype-specific at pre-symptomatic stages and this impact is particularly impactful in female APP/PS1 mice, aligning with our earlier published studies using the 6 Hz kindling protocol (Vande Vyver et al., 2022b).

### APP/PS1 mice are at higher mortality risk after corneal kindling acquisition

While assessing the genotype-related differences in kindling acquisition, we also encountered significant and unanticipated increased premature mortality solely in kindled APP/PS1 mice (Figure 2). Sham kindled APP/PS1 mice were not affected. Our present study directly demonstrates that evoked seizures in APP/PS1 mice, at an age without Aβ accumulation (Figure 8), can alone provoke significant premature mortality (Figure 2B, D). Specifically, this mortality phenotype was remarkable for both males and females after achieving kindling acquisition, with up to a 70% mortality rate. Some, albeit non-significant, mortality was observed during kindling acquisition (Figure 2A, C). There were no notable differences between sexes, suggesting that the mortality is a result of kindling and independent of sex-related differences in seizure threshold. Further corroborating the hypothesis that hyperexcitability preceding aberrant Aβ processing can provoke mortality, we did not observe any significant differences in mortality in both male and female PSEN2-N141I variant versus Tg-control mice (Figure 2E-H). These results reinforce the hypothesis that hyperexcitability in discrete AD-related genotypes directly provokes premature mortality in advance of Aβ accumulation.

**Figure 2.**
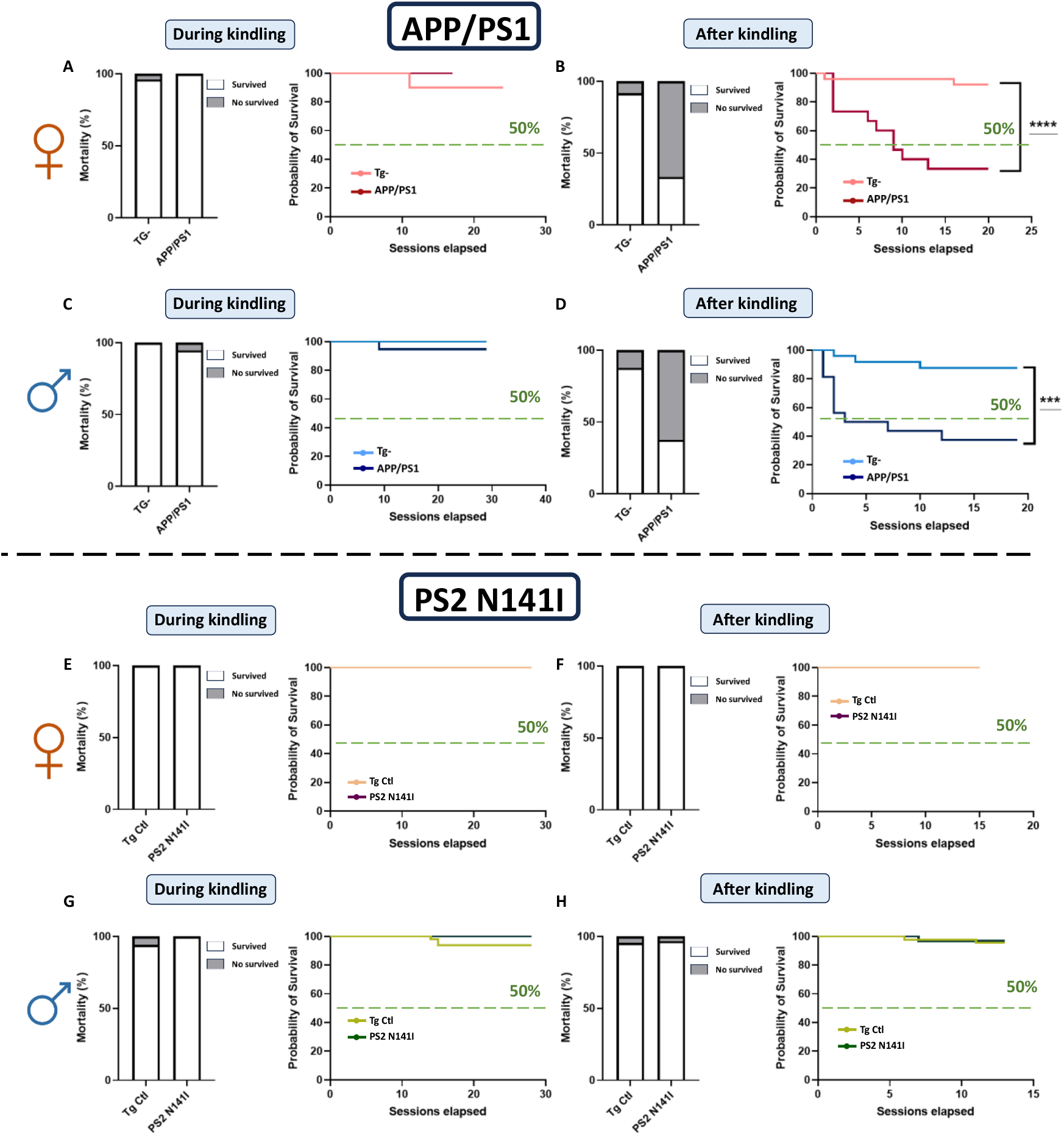
Corneal kindled APP/PS1 mice (2-3 months-old) are highly susceptible to mortality, but survival of non-transgenic controls and PSEN2-N141I transgenic mice is not affected by chronic kindled seizures. **A, C, E, F)** Corneal kindling of 2-3 months-old transgenic (Tg) and non-transgenic mice did not significantly increase mortality during the kindling acquisition period (10-14 days of twice-daily 60 Hz 3 sec 1.6-2.0 mA transcorneal stimulation). However, after animals attained the fully kindled state, both **B)** kindled APP/PS1 females and **D)** APP/PS1 males showed significant and precipitous mortality (≈75%) by 20 sessions. No significant increase in mortality was observed in non-Tg and PSEN2-N141I kindled mice in **F)** females and **H)** males, suggesting that presence of the AD-associated PSEN2 variant does not worsen chronic seizure-induced adverse outcomes. Data for animal survival are presented as a Kaplan-Meier plot and assessed by Log-Rank Mantel Cox test, with * p<0.05 significant. N=15-30 per group. *** indicates p<0.001; **** indicates p<0.0001.

### Chronic evoked seizures are associated with 5-HT pathway dysregulation in young APP/PS1 mice

Considering the extensive and inducible mortality in young APP/PS1 mice subjected to 60 Hz corneal kindling, as well as our earlier similar mortality findings with the 6 Hz kindling model (Vande Vyver et al., 2022a), we hypothesized that this outcome could be linked to changes in molecular pathways associated with mortality risk in epilepsy. The 5-HT pathway may represent a critical component of the pathology present in both epilepsy and AD, and 5-HT system dysfunction is heavily implicated in the pathobiology of SUDEP (Petrucci et al., 2020). To understand the biological underpinnings of premature mortality in kindled APP/PS1 mice, we quantified the protein changes in several relevant components of the 5-HT synthesis and metabolism pathway in HPC and PFC without (sham), and with chronic corneal kindled seizures.

APP/PS1 males and females showed significant downregulation of TPH2 levels in the HPC after they achieved the kindled state compared with kindled Tg- and sham kindled APP/PS1 mice (Figure 3A). Furthermore, MAOA expression was reduced in APP/PS1 kindled mice versus sham-kindled mice (Figure 3A). Surprisingly, SERT protein levels in APP/PS1 mice were overexpressed compared with sham transgenic mice (Figure 3A). No protein changes were appreciable in the 5-HT1A (Figure 3A). Importantly, these findings were not universally detected in AD-associated mice with kindled seizures; PSEN2-N141I mice did not show any hippocampal 5-HT related-protein changes compared with their Tg control (Figure 3B). Thus, 5-HT pathway expression in the HPC showed seizure-induced changes exclusively in APP/PS1 mice.

**Figure 3.**
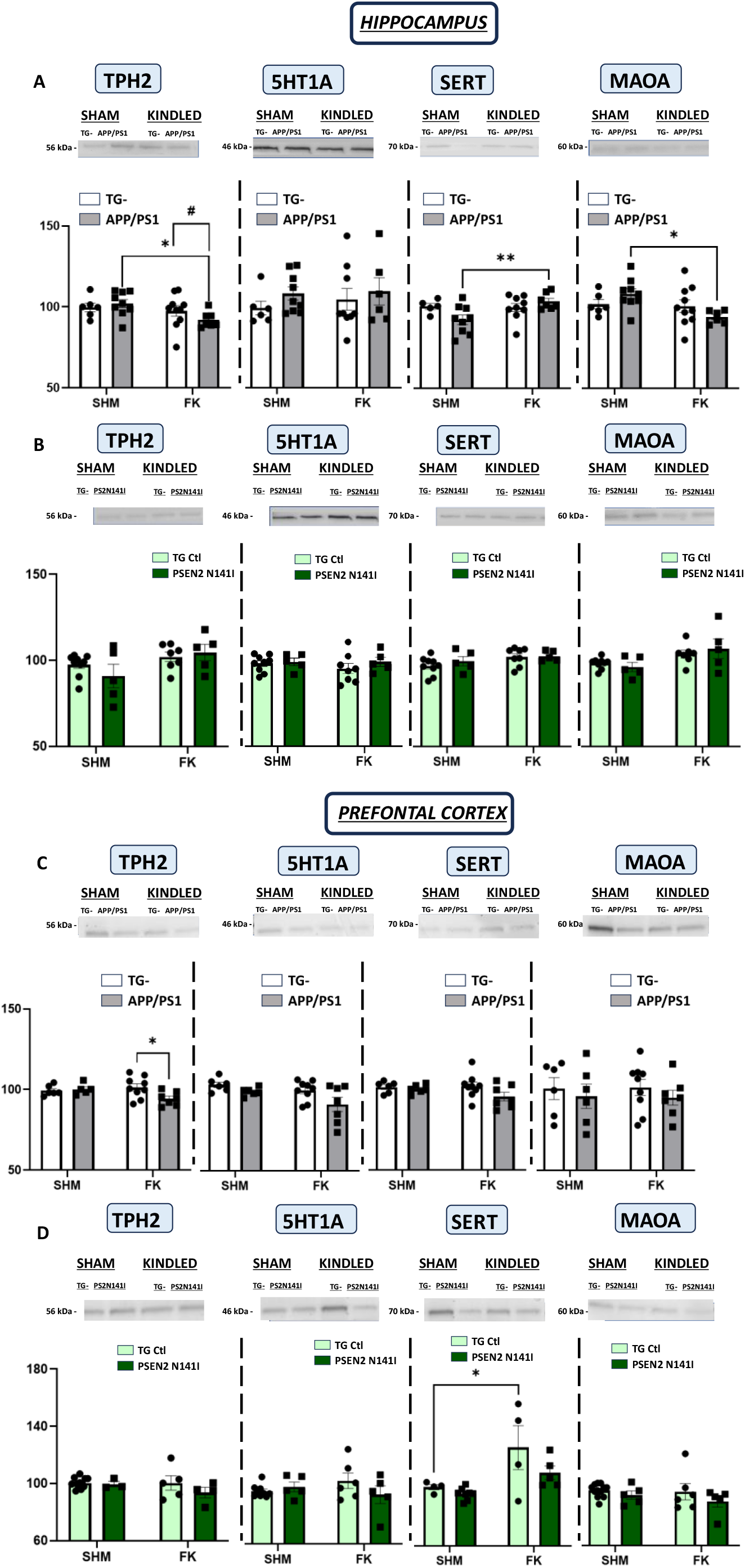
Chronic kindled seizures decrease serotonin synthesis and metabolism protein expression, specifically in isolated hippocampus, in young APP/PS1 mice. **A)** TPH2 and MAOA protein levels in isolated hippocampal homogenates were reduced in kindled male and female APP/PS1 mice relative to their respective genotype-matched SHAM group. Kindled APP/PS1 also exhibited an overexpression in SERT levels compared with SHAM transgenic mice **B)** Protein changes were not observed in kindled or sham PSEN2-N141I mice relative to Tg controls, suggesting that 5-HT synthesis may be reduced solely in APP/PS1 mice with kindled seizures. **C)** TPH2 protein changes were also observed in PFC in APP/PS1 kindled mice. **D)** No further changes were observed in PSEN2 N141I young mice. Bars represented the median and the SEM. Data were assessed by two-way analysis of variance followed by the Bonferroni test (*p < 0.05, **p < 0.01, ***p < 0.005). N= 5-8.

In PFC, APP/PS1 kindled mice showed a reduction in TPH2 levels compared with kindled Tg-mice (Figure 3C). We did not see any changes in protein expression in the remaining 5-HT markers. Like HPC, kindled PSEN2-N141I mice exhibited no changes in serotonergic pathway protein levels in PFC (Figure 3D).

These findings reveal that chronic seizures in the setting of the amyloidogenic model APP/PS1, but prior to the accumulation of Aβ plaques, drives a negative cascade of serotonergic system dysfunction that directly precipitates premature chronic seizure-induced mortality. Importantly, this mortality was not evident in PSEN2-N141I mice nor in sham-kindled APP/PS1 mice, revealing a specific impact of chronic seizures evoked in an APP overexpressing mouse line.

### Chronic kindled seizures accelerate β-amyloid overexpression in the hippocampus of APP/PS1, but non-transgenic controls and PSEN2-N141I mice

Chronic seizures are believed to negatively affect cognitive function and accelerate AD progression (Lehmann et al., 2021). We therefore wanted to assess the impacts of evoked chronic seizures on AD-related protein changes in pre-symptomatic mice of both genotypes to ascertain the degree to which seizures differentially impact AD-associated pathology. Genotype significantly influenced hippocampal protein changes. Although we did not observe significant changes in PS1 expression in any genotype, PS2 protein expression was oppositely impacted in each genotype in a seizure-independent manner (Figure 4A, B). PS2 protein expression was downregulated in APP/PS1 mice independent of kindling status (Figure 4A); conversely PS2 expression was increased in PSEN2-N141I mice regardless of kindling status (Figure 4B). Aβ protein levels in both genotypes resulted in opposite genotype-response. Aβ protein levels were generally elevated in the HPC homogenates of kindled APP/PS1 mice compared with their respective sham and kindled Tg-mice (Figure 4A). PSEN2-N141I mice subjected to corneal kindling did not show any Aβ changes in HPC or PFC by WB analysis (Figure 4B, D). Similar results were observed at APP/PS1 mice (Figure 4C). However, protein changes in PS1 and PS2 were seen at PFC in the APP/PS1 kindled mice (Figure 4C).

**Figure 4.**
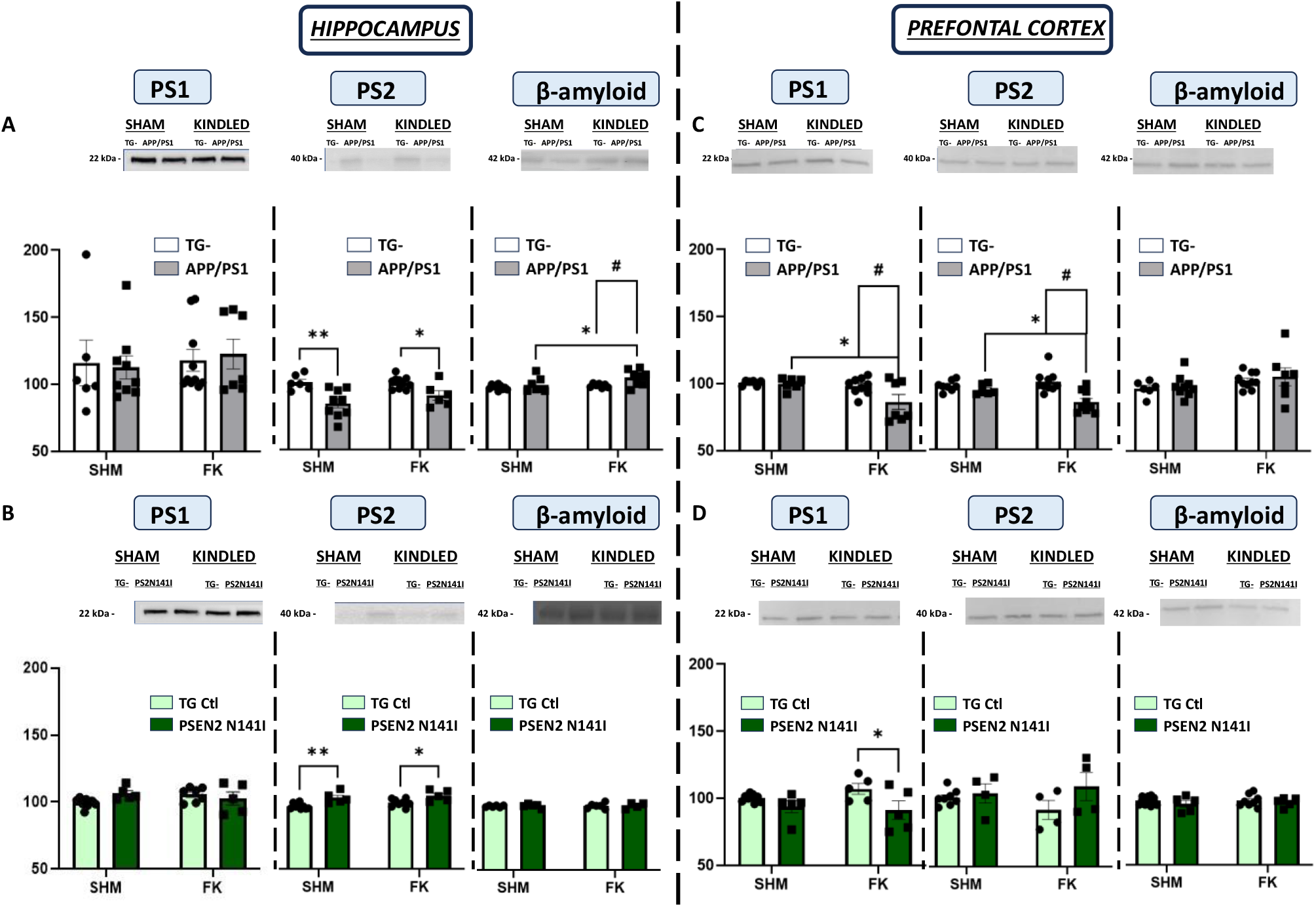
Expression of AD-related proteins in isolated hippocampus are genotype-dependent. PS1 expression in isolated hippocampal homogenates was not altered by kindled seizures in neither **A)** APP/PS1 nor **B)** PSEN2-N141I mouse models of AD relative to their respective control mice at age 2-3-months-old. Protein changes in PS1 were observed in kindled APP/PS1 mice **C)** in PFC, showing a reduction compared with their respective sham littermates and kindled Tg-mice. APP/PS1 mice had reduced expression of the PS2 protein in isolated hippocampal **A)** and PFC **C)** homogenates relative to Tg-in both SHAM and kindled mice, suggesting that corneal kindled seizures do not alone modify expression of the PS2 protein at this age in mice of the APP/PS1 genotype. Conversely, hippocampal PS2 protein expression was generally increased in PSEN2-NI141 mice relative to their matched Tg control **B)**, and this expression was not affected by kindled seizure history. **A)** Aβ protein overexpression was detected in APP/PS1 kindled young mice relative to their matched Tg- and sham APP/PS1 littermates. **B)** PSEN2 N141I mice did not exhibit any changes in Aβ protein levels. There was no kindling-induced change in Aβ protein expression in prefrontal cortex homogenates in neither **C)** APP/PS1 nor **D)** PSEN2-N141I mice relative to their matched controls. Bars represented the median and the SEM. Data were assessed by two-way analysis of variance followed by the Bonferroni test (*p < 0.05, **p < 0.01, ***p < 0.005). N= 5-8.

These findings suggest that chronic evoked seizures alter AD-related protein expression in a genotype-dependent manner in young, pre-symptomatic AD mice. There are genotype-specific changes in PS2 protein expression, suggesting that overexpression of this protein may protect against neuronal hyperexcitability which warrants further detailed investigation. Moreover, kindling-induced upregulation of Aβ protein levels in HPC in the absence of overt plaque accumulation suggests that chronic neuronal hyperexcitability can directly accelerate AD-pathology in advance of the identification of AD-hallmarks. Importantly, these findings demonstrate that unchecked neuronal hyperexcitability is a specific driver of worsened functional and pathological outcomes in AD-associated mouse models.

### APP/PS1 mice exhibit a decoupling of the serotonin pathway protein expression

The 5-HT pathway has been proposed to be involved in the pathology of epilepsy and AD and is heavily implicated in SUDEP pathology (Buchanan et al., 2014). Only kindled APP/PS1 mice exhibited changes in 5-HT synthesis (TPH2) and metabolism (MAOA) markers in isolated HPC. Moreover, kindled APP/PS1 mice also showed reduced PS2 expression and increased Aβ protein levels. These protein differences suggest a seizure-induced decoupling of the serotonin signaling pathway that drives AD-related protein modifications. To confirm this, we assessed the potential correlation between the AD and serotonin markers in HPC (Figure 5). APP/PS1 sham and kindled young mice demonstrated important correlations between the targets that we studied. Interestingly, hippocampal MAOA expression vs hippocampal Aβ protein levels were significantly negatively correlated in kindled APP/PS1 mice (Figure 5E), suggesting that 5-HT signaling is a critical contributor to AD-pathological hallmark progression. No correlation was evident in sham kindled APP/PS1 mice (Figure 5E) or PSEN2-N141I kindled or sham mice (not shown). Furthermore, sham-kindled APP/PS1 mice showed a positive correlation between PS2 and SERT (Figure 5F). This correlation was lost in kindled APP/PS1 mice. No remarkable findings were observed in PSEN2-N141I mice (Supplemental Figure 2). These results suggest that at pre-symptomatic stages, 2-month-old kindled APP/PS1 mice exhibit a decoupling between 5-HT synthesis and metabolism that likely accelerates the deposition of AD-related pathological hallmarks.

**Figure 5:**
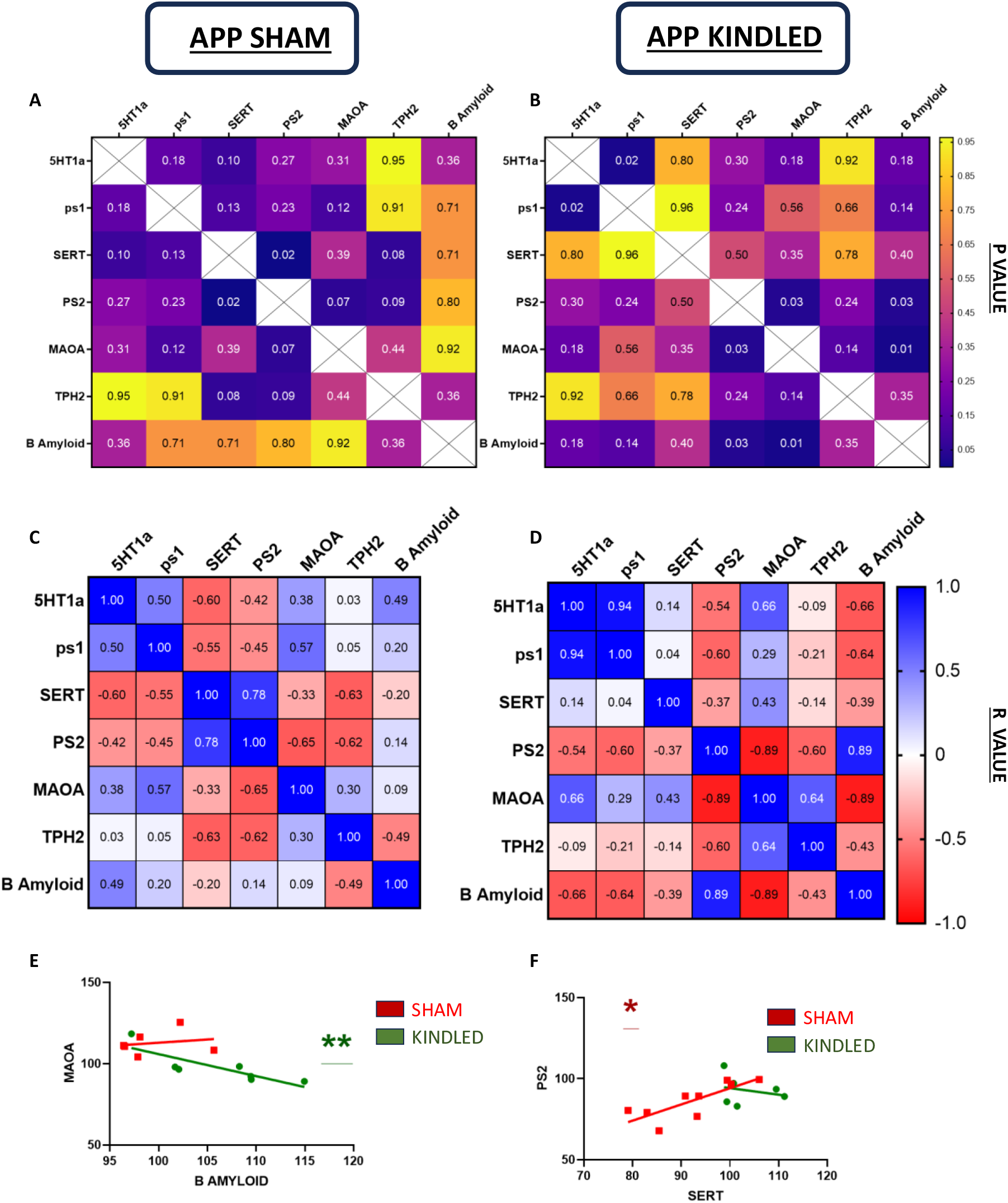
5-HT metabolism and transport enzymes, Aβ, and PS2 expression are tightly correlated in kindled APP/PS1 mice. Correlational analysis was assessed in both sham and kindled APP/PS1 mice to study the hippocampal interaction between AD-related proteins and 5-HT pathway enzyme levels. Figures A and B show the *p value* obtained when two protein levels were measured in APP/PS1 sham and kindled mice, respectively. Figures C and D exhibit the *R value* obtained in both sham and kindled APP/PS1 mice, respectively, to determine if the correlation is inverse or straight. **E)** Comparison between hippocampal MAOA and Aβ protein levels revealed an inverse correlation in the kindled APP/PS1 mice but not in the sham mice, demonstrating the connection between 5-HT and AD pathology in the context of seizures. **F)** Sham APP/PS1 mice exhibited a positive correlation between PS2 and SERT levels, which was not evident in APP/PS1 kindled mice, demonstrating the decoupling of 5-HT and AD-related proteins under chronic evoked seizures. Correlational studies were performed following Spearman correlation. *p < 0.05.

### Chronic evoked seizures promote an increase in glial response in young APP/PS1 mice independent of Aβ plaque accumulation

Astrocytes have been demonstrated to be a key player regulating hyperexcitability and glutamate release in epilepsy (Barker-Haliski & White, 2015). Furthermore, activated microglia and neuroinflammation is involved in AD pathology (Revuelta et al., 2022), including Aβ deposition (Turkin et al., 2021). Enhancing the 5-HT tone is a promising strategy to counteract the inflammatory-related mechanisms in AD, including Aβ clearance (Turkin et al., 2021). Since both the APP/PS1 and PSEN2-N141I mouse models are associated with altered microglia reactivity (Cao et al., 2021; Fung et al., 2020), we evaluated if the seizure-induced changes in 5-HT pathway protein expression similarly resulted in glial dysfunction and Aβ accumulation at pre-symptomatic stages. To perform this analysis, we assessed the intensity of signal of astrocytes (GFAP), microglia (Iba-1), and Aβ (6e10/Thioflavin) in HPC (DG, CA1, CA3) and CTX.

In line with our earlier studies (Knox KM, 2023; Loewen et al., 2016), corneal kindling increase GFAP immunoreactivity in HPC structures (*p: 0.27; F: 5.163*) (Figure 6). However, significant differences were only observed in kindled APP/PS1 mice in DG (*p=0.028*), and CA3 (*p=0.023*; Figure 6B) when compared to Tg-sham-kindled mice. Iba-1 reactivity exhibited similar results. Young APP/PS1 mice showed an increase in Iba-1 levels in DG, CA1 and CA3 compared with Tg-sham young mice (Figure 7B). In this case, no significant differences were observed in CTX overlaying dorsal CA1. Interestingly, these inflammatory changes did not trigger any Aβ plaque accumulation; neither the sham nor the kindled APP/PS1 mice demonstrated evidence of Aβ plaque deposition in HPC (Figure 8). To confirm our labeling was specifically capable of detecting Aβ plaques, aged (+10 months-old), kindled APP/PS1 mice (Figure 8) were included confirming the specificity of our labeling technique. No remarkable protein changes in GFAP (Figure 6C), Iba-1 (Figure 7C) or 6e10 (not shown) were observed in young PSEN2-N141I sham or kindled mice. Additionally, NeuN staining showed no remarkable differences between either of the groups confirming that corneal kindling in the presence of and AD-related mutations does not trigger neurodegeneration at early stages (Figure Supplementary 3). These findings altogether demonstrate that chronic evoked seizures induce pathological changes in the setting of Aβ overexpression in an amyloidogenic mouse model, which could cumulatively accelerate functional decline AD in the absence of overt Aβ plaques deposition.

**Figure 6:**
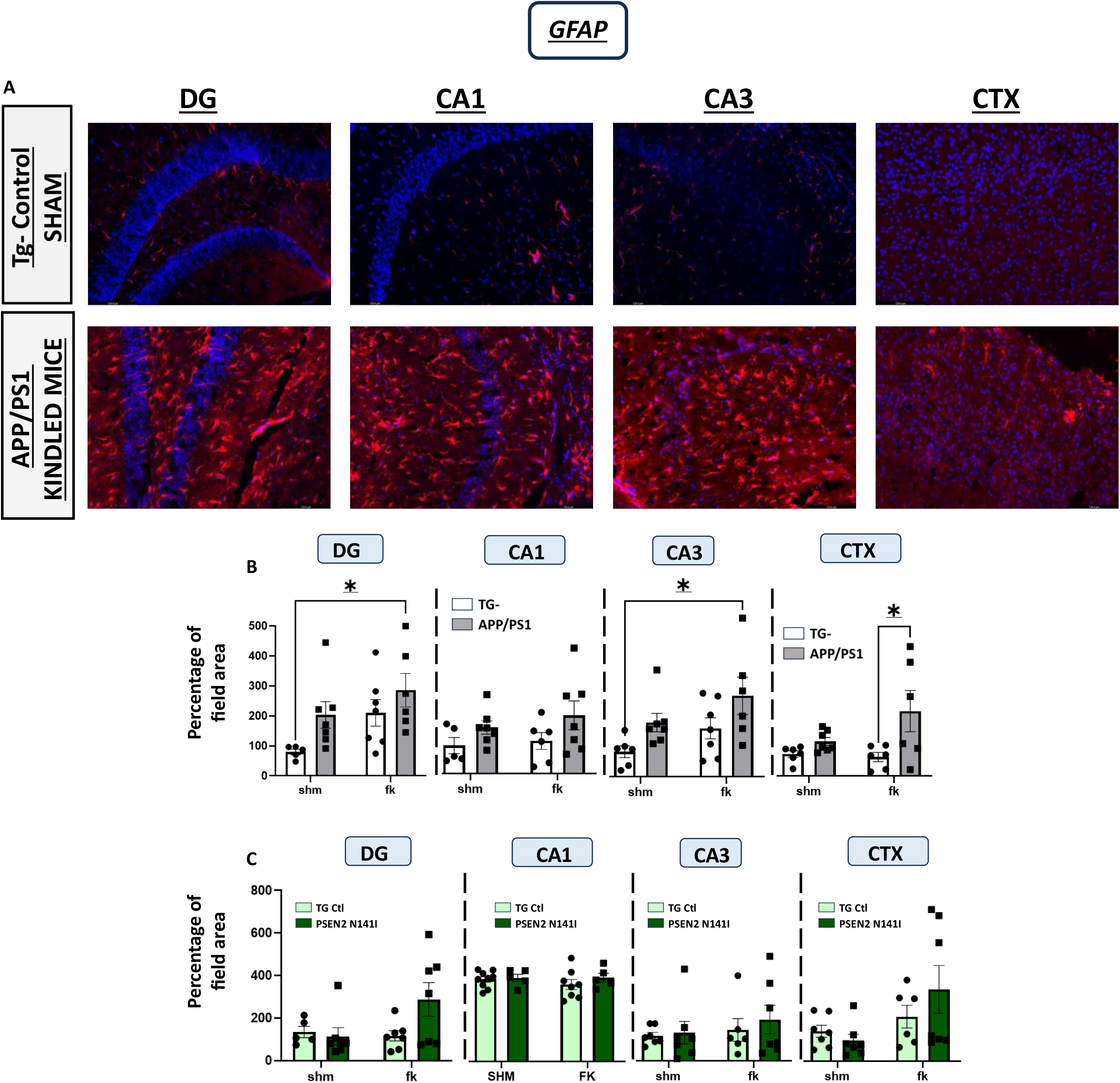
Chronic kindled seizures evoke increased GFAP expression in posterior parietal association cortex and dorsal hippocampus of APP/PS1 mice. **A)** Representative images of DAPI+GFAP staining at DG, CA1, CA3 and CTX (posterior parietal association cortex overlays area CA1) of Tg-sham and APP/PS1 kindled mice. **B)** Histological studies demonstrated that under the influence of chronic seizures, APP/PS1 young mice exhibited an increase in GFAP reactivity in DG, CA3 and CTX compared with the controls. **C)** PSEN2 N141I mice did not show any significant increase in GFAP levels compared with their Tg controls. Overall, these results confirm that kindling triggers a pathological increase in astrogliosis. Bars represent the median and the SEM. Data were assessed by two-way analysis of variance followed by the Bonferroni test (*p < 0.05, **p < 0.01, ***p < 0.005). N= 5-8.

**Figure 7:**
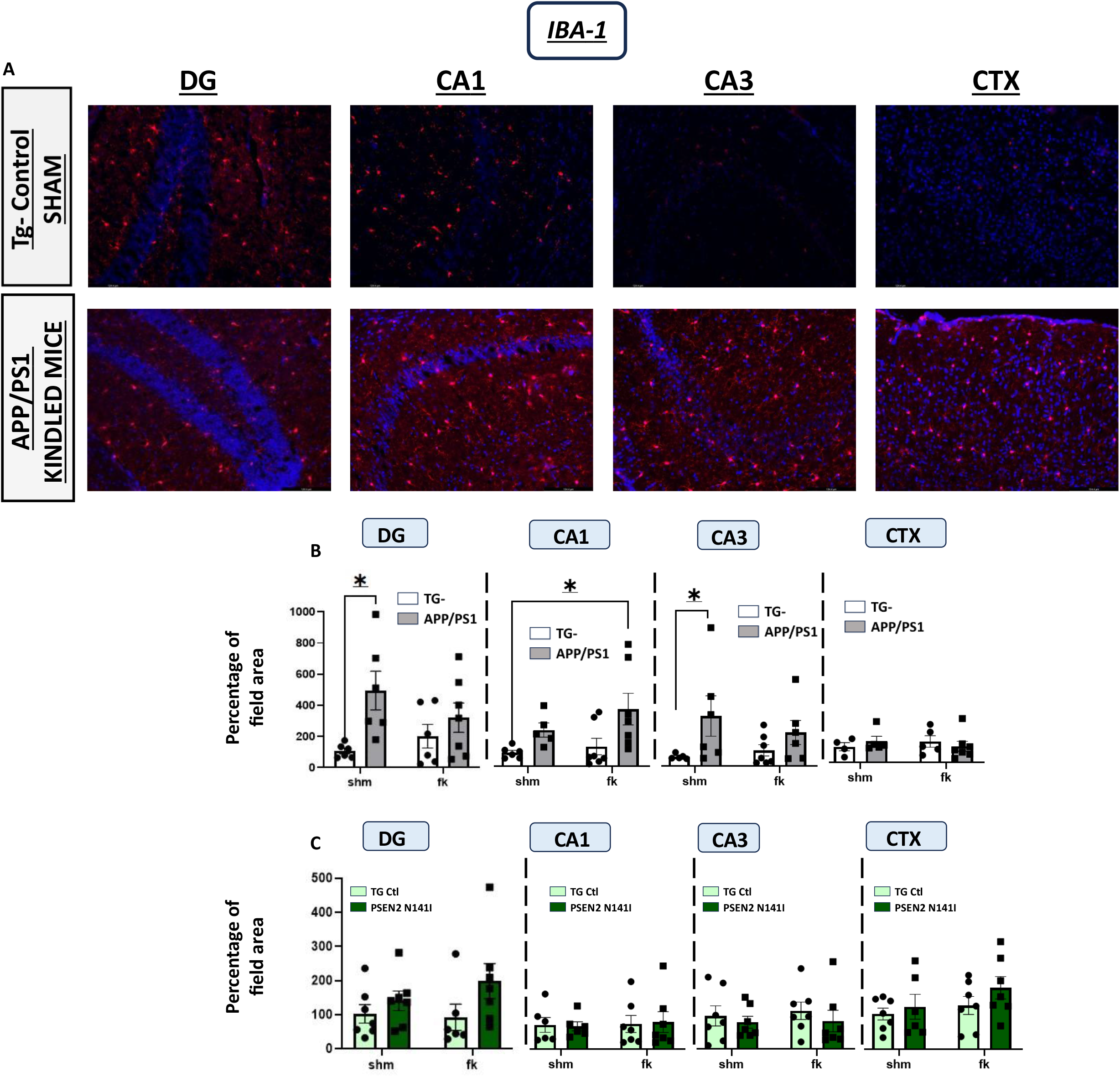
Microgliosis is increased in cortical and hippocampal brain regions in corneal kindled APP/PS1 mice. **A)** Representative images of DAPI+Iba1 staining at HPC and CTX of Tg-sham and APP/PS1 kindled mice. **B)** Histological studies demonstrate APP/PS1 mice subjeted to corneal kindling exhibit an increase in hippocampal Iba1 reactivity, a marker of activated microglia. **C)** PSEN2-N141I mice did not show any significant increase in Iba1 immunoreactivity. Bars represent the median and the SEM. Data were assessed by two-way analysis of variance followed by the Bonferroni test (*p < 0.05, **p < 0.01, ***p < 0.005). N= 5-8.

**Figure 8:**
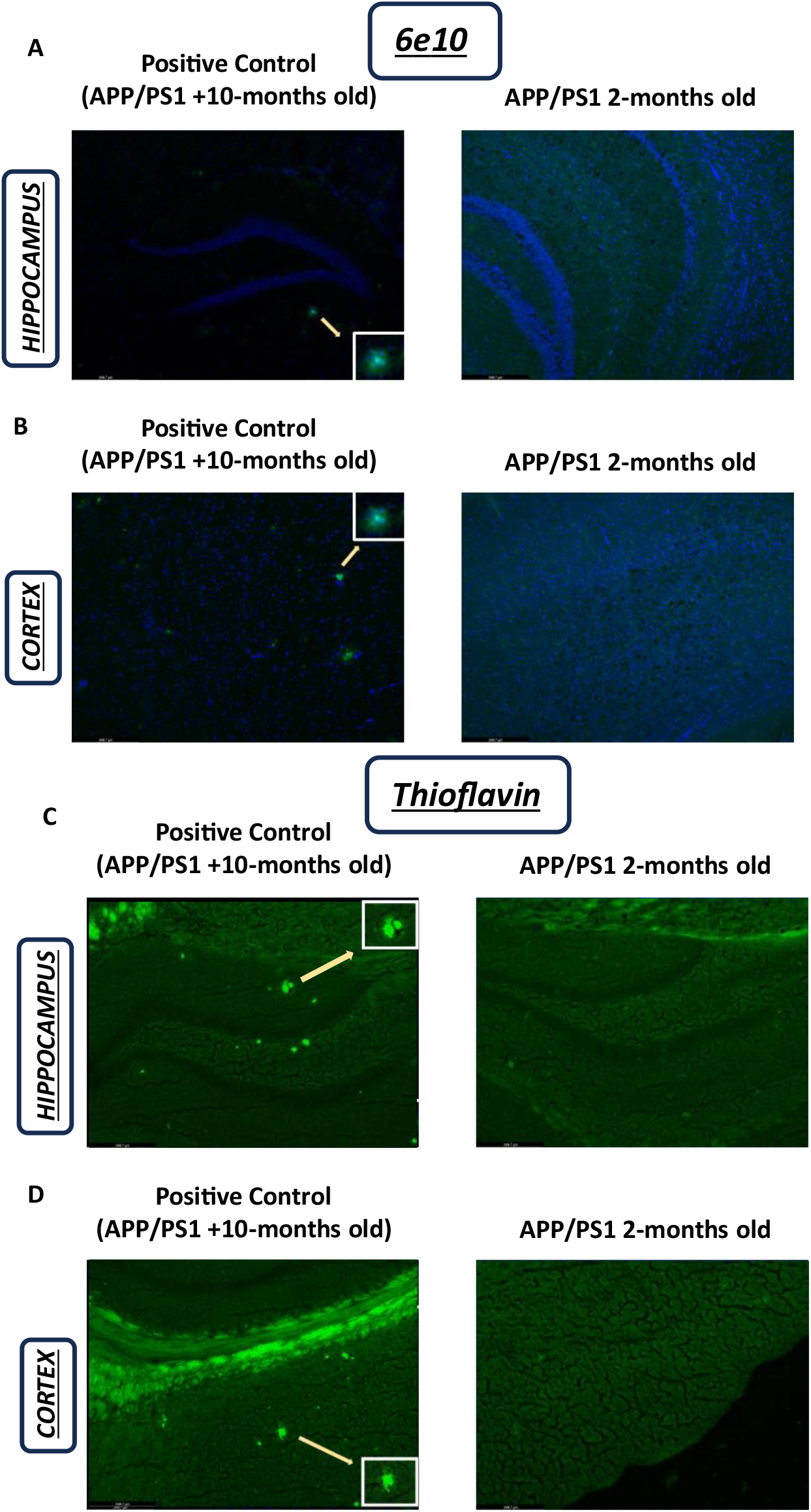
There is no evidence of Aβ plaque accumulation corneal kindled young APP/PS1 mice. **A)** Representative photomicrographs of 6e10 staining in HPC of >10 months-old APP/PS1 mouse (positive control) and a young APP/PS1 mouse. **B)** Representative images of 6e10 staining in CTX of >10 months-old APP/PS1 mouse (positive control) and a young APP/PS1 mouse. **C)** Representative photomicrographs of Thioflavine staining in HPC of >10 months-old APP/PS1 mouse (positive control) and a young APP/PS1 mouse. **D)** Representative images of Thioflavin staining in CTX of >10 months-old APP/PS1 mouse (positive control) and a young APP/PS1 mouse. Overall, these results demonstrate that the pathological changes observed in corneal kindled APP/PS1 mice occurs at pre-symptomatic stages.

## Discussion

While seizures can present in people with AD, the causative mechanisms are still relatively unknown. It is likely that untreated focal and myoclonic seizures negatively affect the cognitive and pathological burden of AD, including increased risk of mortality (Lehmann et al., 2021; Zawar, 2022). Our present study provides unequivocal direct evidence that chronic seizures negatively affect survival in mice with specific AD-associated risk factors. In this work, we also evaluated the potential mechanisms that explain the susceptibility to seizures and their sequelae in two murine models of EOAD at an age well-before Aβ plaque deposition (Minkeviciene et al., 2008). APP/PS1 mice were highly susceptible to 60 Hz corneal kindling and mortality versus Tg-littermate mice. These effects were not observed in PSEN2-N141I mice when compared to their matched Tg controls. These present findings are reminiscent of earlier work demonstrating greater susceptibility of young APP/PS1 and 3xTg mice to 6 Hz kindling (Vande Vyver et al., 2022b) and of aged Tg2576 mice to amygdala kindling (Chan et al., 2015), but further illustrates stark differences in kindling susceptibility of mice with PSEN2 variants or loss of normal function (Beckman et al., 2020). Further, we herein illustrate that expression of an AD-associated genotype alone does not drive seizure-induced excess mortality and that this mortality can arise well-before Aβ plaques accumulate. Interestingly, our present study demonstrates that pre-symptomatic APP/PS1 kindled mice exhibit changes in the expression of 5-HT synthesis and degradation enzymes exclusively in the isolated HPC and PFC, suggesting possible deficits in 5-HT release as a result. These two AD models showed innate differences in the expression of the PS2 protein, revealing a potentially novel modifier of vulnerability to seizures and formation of a hyperexcitable neuronal network that warrants further study. APP/PS1 kindled mice also exhibited an exacerbated glial response suggesting a vicious cycle between 5-HT, inflammation, and AD-related changes that occur prior to pathological symptomatic stages of AD. These modifications resulted in the observed hippocampal increase of β-amyloid protein without overt plaques accumulation. Altogether, our present study reveals substantial differences in the functional and molecular impacts of kindled seizures in mice with AD-associated genotypes at an age prior to overt Aβ pathology. This study emphasizes the urgency in carefully considering pharmacotherapeutic management of seizures in AD as an addressable drug repurposing opportunity to reduce the burden of AD.

We herein demonstrate extensive seizure-induced mortality coincident with dysregulation in 5-HT pathway protein expression that arises exclusively in APP/PS1 mice. 5-HT is one of the most promising targets driving SUDEP risk in epilepsy (Buchanan et al., 2014; DeGiorgio et al., 2019; Feng & Faingold, 2017; Massey et al., 2021; Petrucci et al., 2020; Richerson & Buchanan, 2011; Tupal & Faingold, 2021). Additionally, mice lacking the 5HT2C receptor have high mortality (Massey et al., 2021) and 5-HT agonists, such as fenfluramine, may prevent seizure-induced preclinical and clinical mortality (Cross et al., 2021; Zhang et al., 2015). Furthermore, AD also promotes 5-HT pathway dysregulation, possibly due to excessive neuronal hyperexcitability (von Linstow et al., 2022). As in AD patients, spontaneous seizures occur in APP/PS1 mice (DeGiorgio et al., 2020; Reyes-Marin & Nunez, 2017). Emerging clinical evidence would also suggest that people with AD and comorbid seizures are at increased mortality risk relative to individuals without seizures in AD (DeGiorgio et al., 2020; Zawar, 2022). Unlike prior preclinical studies of spontaneous recurrent seizures in APP/PS1 mice, ours is the first to demonstrate overt and inducible mortality before Aβ deposition as a direct result of chronic 60 Hz seizures, in mice aged equivalent to a ∼25-year-old human. 5-HT imbalance has been also reported in old APP/PS1 mice at symptomatic stages (Lin et al., 2015) and APP/PS1 mice lacking the TPH2 enzyme also demonstrate heightened mortality (von Linstow et al., 2022). Tph2 inactivation influences APP processing, at least in the HPC (von Linstow et al., 2022), showing changes in Aβ protein levels like we observed. However, seizure-induced 5-HT pathway disruptions at pre-symptomatic stages have never been evaluated in AD-associated models. We found that chronic seizures reduce TPH2 expression and increase mortality exclusively in kindled APP/PS1 mice. TPH2 downregulation secondary to seizures has been previously reported in other epilepsy models with heightened mortality risk (Chen et al., 2019). Hippocampal MAOA protein expression was similarly affected by kindling only in APP/PS1 mice. MAOA downregulation may reflect a compensatory effect secondary to the reduction in 5-HT levels or seizure-induced inflammation leading to reduced MAOA levels (Beroule, 2020). Further, MAOA expression correlated to Aβ protein levels only in APP/PS1 kindled mice, indicating the 5-HT seizure-induced changes can trigger some undiagnosed AD mechanisms at pre-symptomatic stages. Our data cumulatively indicate that the combination of APP/PS1 genotype and chronic seizures leads to significant reductions in 5-HT synthesis and metabolism enzymes prior to pathological Aβ deposition, pointing to a conserved role for seizures eliciting deficits in 5-HT release as a plausible mechanism underlying the seizure-associated mortality in young APP/PS1 mice. More importantly, these findings suggest a possible novel non-canonical SUDEP model that carries significant potential for addressing the most catastrophic outcome of uncontrolled epilepsy.

Considering that PSEN2-N141I mice were largely protected from chronic seizure-induced mortality and disruptions in 5-HT synthesis and metabolism enzyme expression, another important novel aspect of this work is that PS2 overexpression may differentially influence susceptibility to neuronal hyperexcitability and accordingly, seizure-induced excess mortality. Indeed, PS2 protein expression was reduced in APP/PS1 mice versus Tg-controls, whereas it was increased in PSEN2-N141I mice relative to their Tg control mice, suggesting a critical contribution of this protein to seizure susceptibility that warrants further study. Neuronal hyperexcitability driven by disrupted calcium handling is a crucial factor in epilepsy and AD (Dibue et al., 2013; Ge et al., 2022). Indeed, the antiseizure medicine levetiracetam, which alleviates cognitive decline in animal models and in patients with AD and epileptiform activity, exerts its anticonvulsant effect and preserves neuronal viability by restoring calcium homeostasis (Pisani et al., 2004; Vossel et al., 2021; Zheng et al., 2022). How PS proteins modulate calcium homeostasis is still largely unknown. Some studies suggest that PS1 and PS2 variations increase intracellular calcium, resulting in an increase in hyperexcitability and neurodegeneration at pre-symptomatic stages (Galla et al., 2020). Moreover, animals with PSEN2 variants also have reduced calcium levels (Galla et al., 2020), suggesting inconclusive information about this phenomenon. Our data suggests that PSEN2-N141I and PSEN1-dE9 mutations trigger opposite presenilin-related mechanisms that exert different effects on seizure vulnerability, which requires further investigation. We herein demonstrate that increased expression of PS2 protein may be anticonvulsant in young animals, as evidenced by reduced seizure susceptibility in PSEN2-N141I mice (Figure 1). The PSEN2-N141I variant may be influencing the seizure-induced disruptions in calcium homeostasis that is commonly observed in the hyperexcitability of epilepsy. On the other hand, APP/PS1 mice demonstrated reduced PS2 protein expression, possibly translating to excess accumulation of intracellular calcium that ultimately increases seizure susceptibility in AD.

It is essential to note that AD is a complex and multifactorial condition, and the interactions between various cellular and molecular players, including PS2 and microglia, are still an area of ongoing research. An increase in the expression of GFAP and Iba-1 is commonly considered a hallmark of neuroinflammation in both AD (Fung et al., 2020) and epilepsy (Eyo et al., 2017). In fact, astrocytes, together with microglia, react to a diverse range of inflammatory agents, including Aβ plaque deposition (Gonzalez-Reyes et al., 2017). Since none of the genotypes in our study exhibited any Aβ plaques, other non-amyloidogenic mechanisms may be involved in this glial response. It is likely that 5-HT plays a pivotal role in AD and epilepsy pathology. Notably, 5-HT also modulates proliferation, development, and migration of microglia (Albertini et al., 2023). Microglia also express the Htr2b gene, which encodes the 5-HT2B receptor (Kolodziejczak et al., 2015; Krabbe et al., 2012). In the lifelong absence of microglial 5-HT2B receptor, peripheral LPS injection causes cytokine overexpression and prolonged neuroinflammation *in vivo* that represents an increased pro-inflammatory phenotype (Turkin et al., 2021). These glial and 5-HT changes in APP/PS1 kindled mice may bidirectionally affect PS2 protein changes. It is not well understood how 5-HT metabolism and PS2 can interact; however, we know that specific loss of PS2 expression results in an exaggerated cytokine response by microglia (Jayadev, Case, et al., 2010). Seizure-induced decoupling between PS2 and 5-HT was also confirmed with the correlational studies performed in APP/PS1 kindled mice (Figure 5). The pivotal role of PS2 in the regulation of 5-HT and Iba-1 was corroborated in the PSEN2-N141I kindled mice, who exhibited an upregulation of PS2 protein resulting in no significant differences in GFAP, Iba-1 and 5-HT protein levels compared to their respective controls. Additionally, NeuN staining in HPC (Supplementary Figure 3) showed no significant differences in the number of neurons between groups, confirming that hippocampal neurodegeneration did not contribute to any of the observed differences in survival, seizure risk, or biomolecular changes otherwise observed. Hence, dysregulation of PS protein expression in AD may induce other pathological changes independent of Aβ accumulation, such as changes in 5-HT synthesis or metabolism pathways, resulting in the consequent increased neuroinflammation.

AD drug development remains susceptible to lurking problems capable of compromising future research efforts (Mehta et al., 2017). Anti-Aβ monoclonal antibodies (mAβs) have dominated the therapeutic development landscape for decades, with only a handful of agents demonstrating small, but significant, biologically-meaningful benefit to slow or steady the course of AD (Budd Haeberlein et al., 2022; van Dyck et al., 2022). Hundreds of compounds that have been tested as potential therapies were either abandoned in development or failed in clinical trials (Mehta et al., 2017). Aβ is appealing as a therapeutic target as it is extra-neuronal and is associated with toxicity to the milieu (Mehta et al., 2017). However, Aβ load does not necessarily correlate with cognitive decline. Resilience to AD in the face of high Aβ pathology and neurofibrillary tangles is increasingly challenging the traditional Aβ hypothesis of AD (Arenaza-Urquijo & Vemuri, 2018; Lopera et al., 2023), potentially offering an explanation as to why administration of mAβs in symptomatic stages of disease does not necessarily translate into clinically meaningful improvements in disease burden. Instead, these clinical failures may suggest that the treatment of symptomatic AD occurs too late in the disease course (Mehta et al., 2017). Our present study provides critical direct evidence that excess neuronal hyperexcitability leading to 5-HT dysfunction and attendant neuroinflammation may instead incite the onset of AD pathology (i.e., increased Aβ levels and neuroinflammation) such that plaque deposition is a functional consequence of this hyperexcitability-initiated pathological cascade. In this sense, uncovering the mechanisms that trigger hyperexcitability and the resulting Aβ plaque deposition, pathological decline, and excess mortality may represent a more promising strategy to modify the course of AD. Managing seizures and hyperexcitability in EOAD thus represents a tractable drug repurposing avenue to minimize functional decline and reduce risk of excess mortality in people with AD.

## Conclusions

Our present study reveals several critical novel findings of relevance to the study of seizures in EOAD, as well as presents critical evidence that neuronal hyperexcitability in advance of canonical Aβ accumulation accelerates functional decline at pre-symptomatic stages (Supplemental Figure 1B). First, we illustrate substantial differences in seizure susceptibility in discrete AD-associated mouse models at young, pre-symptomatic ages. This provides further supporting evidence of the heterogeneity in AD-related genotypes and associated disease course. Second, we demonstrate marked differences in the mortality of APP/PS1 mice secondary to evoked seizures, confirming the deleterious effect of seizures in AD well-prior to pathologic stages of disease. This work unequivocally demonstrates that seizures, and not Aβ plaque accumulation, initiates a negative cascade to accelerate mortality in an AD model. Third, we report that chronic seizures elicit marked depletion of TPH2 and MAOA enzyme levels in APP/PS1, but not PSEN2-N141I, mice, suggesting that seizure-induced mortality and overexpression of Aβ protein in the APP/PS1 model is due to deficits in 5-HT release. Fourth, PS1:PS2 imbalance pivotally impacts glial modulation and seizure protection. Considering the failure of historical AD therapeutic strategies, these data cumulatively suggest that untreated chronic seizures may accelerate the pathological mechanisms involved in the development of AD in advance of Aβ plaque accumulation. Prior studies of seizures in AD mouse models had suggested that seizures secondary to Aβ accumulation were responsible for the precipitous declines in survival in APP/PS1 mice (von Linstow et al., 2022). However, our present study now clearly demonstrates that seizures, and not Aβ accumulation *per se*, are the driving factor underlying excess mortality in this mouse model and that deficits in 5-HT release and glial reactivity may be culpable. Future studies are needed to further characterize how inflammatory response triggers seizure-induced AD onset to further define whether it represents a novel strategy to intervene at pre-symptomatic stages of the disease. Overall, those studies would be critical to identify novel pathological features of AD independent of AD-hallmarks, such as the presence of seizures or 5-HT dysregulation.

## Acknowledgements

American Epilepsy Society Junior Investigator Award and NIA (R01AG067788) to MBH.

## Details mentioning the exact contribution of each author to the work submitted

A.D.P.: conceptualization, methodology, investigation, data curation, writing—original draft. L.R., K.M.K., L.L.: investigation, data curation, writing—original draft. S.D: investigation, writing— original draft; S.J.: resources, writing—review and editing; M.B.H.: conceptualization, methodology, investigation, formal analysis, funding acquisition, supervision, resources, writing— original draft, writing—review and editing.

## Conflict of Interest/Ethical Publication Statement

None of the authors has any conflict of interest to disclose. We confirm that we have read the Journal’s position on issues involved in ethical publication and affirm that this report is consistent with those guidelines.

## References

Abbink, M. R., Kotah, J. M., Hoeijmakers, L., Mak, A., Yvon-Durocher, G., van der Gaag, B., Lucassen, P. J., & Korosi, A. (2020). Characterization of astrocytes throughout life in wildtype and APP/PS1 mice after early-life stress exposure. J Neuroinflammation, 17(1), 91. https://doi.org/10.1186/s12974-020-01762-z

Albertini, G., D’Andrea, I., Druart, M., Bechade, C., Nieves-Rivera, N., Etienne, F., Le Magueresse, C., Rebsam, A., Heck, N., Maroteaux, L., & Roumier, A. (2023). Serotonin sensing by microglia conditions the proper development of neuronal circuits and of social and adaptive skills. Mol Psychiatry. https://doi.org/10.1038/s41380-023-02048-5

Arenaza-Urquijo, E. M., & Vemuri, P. (2018). Resistance vs resilience to Alzheimer disease: Clarifying terminology for preclinical studies. Neurology, 90(15), 695–703. https://doi.org/10.1212/WNL.0000000000005303

Barker-Haliski, M., & White, H. S. (2015). Glutamatergic Mechanisms Associated with Seizures and Epilepsy. Cold Spring Harb Perspect Med, 5(8), a022863. https://doi.org/10.1101/cshperspect.a022863

Barker-Haliski, M. L., Vanegas, F., Mau, M. J., Underwood, T. K., & White, H. S. (2016). Acute cognitive impact of antiseizure drugs in naive rodents and corneal-kindled mice. Epilepsia, 57(9), 1386–1397. https://doi.org/10.1111/epi.13476

Beckman, M., Knox, K., Koneval, Z., Smith, C., Jayadev, S., & Barker-Haliski, M. (2020). Loss of presenilin 2 age-dependently alters susceptibility to acute seizures and kindling acquisition. Neurobiol Dis, 136, 104719. https://doi.org/10.1016/j.nbd.2019.104719

Beroule, D. G. (2020). Paradoxical Effects of a Cytokine and an Anticonvulsant Strengthen the Epigenetic/Enzymatic Avenue for Autism Research. Front Cell Neurosci, 14, 585395. https://doi.org/10.3389/fncel.2020.585395

Buchanan, G. F., Murray, N. M., Hajek, M. A., & Richerson, G. B. (2014). Serotonin neurones have anti-convulsant effects and reduce seizure-induced mortality. J Physiol, 592(19), 4395–4410. https://doi.org/10.1113/jphysiol.2014.277574

Budd Haeberlein, S., Aisen, P. S., Barkhof, F., Chalkias, S., Chen, T., Cohen, S., Dent, G., Hansson, O., Harrison, K., von Hehn, C., Iwatsubo, T., Mallinckrodt, C., Mummery, C. J., Muralidharan, K. K., Nestorov, I., Nisenbaum, L., Rajagovindan, R., Skordos, L., Tian, Y., Sandrock, A. (2022). Two Randomized Phase 3 Studies of Aducanumab in Early Alzheimer’s Disease. J Prev Alzheimers Dis, 9(2), 197–210. https://doi.org/10.14283/jpad.2022.30

Cao, S., Fisher, D. W., Rodriguez, G., Yu, T., & Dong, H. (2021). Comparisons of neuroinflammation, microglial activation, and degeneration of the locus coeruleus-norepinephrine system in APP/PS1 and aging mice. J Neuroinflammation, 18(1), 10. https://doi.org/10.1186/s12974-020-02054-2

Chan, J., Jones, N. C., Bush, A. I., O’Brien, T. J., & Kwan, P. (2015). A mouse model of Alzheimer’s disease displays increased susceptibility to kindling and seizure-associated death. Epilepsia, 56(6), e73–77. https://doi.org/10.1111/epi.12993

Chen, Q., Tian, F., Yue, Q., Zhan, Q., Wang, M., Xiao, B., & Zeng, C. (2019). Decreased serotonin synthesis is involved in seizure-induced respiratory arrest in DBA/1 mice. Neuroreport, 30(12), 842–846. https://doi.org/10.1097/WNR.0000000000001287

Cross, J. H., Galer, B. S., Gil-Nagel, A., Devinsky, O., Ceulemans, B., Lagae, L., Schoonjans, A. S., Donner, E., Wirrell, E., Kothare, S., Agarwal, A., Lock, M., & Gammaitoni, A. R. (2021). Impact of fenfluramine on the expected SUDEP mortality rates in patients with Dravet syndrome. Seizure, 93, 154–159. https://doi.org/10.1016/j.seizure.2021.10.024

DeGiorgio, C. M., Curtis, A., Carapetian, A., Hovsepian, D., Krishnadasan, A., & Markovic, D. (2020). Why are epilepsy mortality rates rising in the United States? A population-based multiple cause-of-death study. BMJ Open, 10(8), e035767. https://doi.org/10.1136/bmjopen-2019-035767

DeGiorgio, C. M., Curtis, A., Hertling, D., & Moseley, B. D. (2019). Sudden unexpected death in epilepsy: Risk factors, biomarkers, and prevention. Acta Neurol Scand, 139(3), 220–230. https://doi.org/10.1111/ane.13049

Dibue, M., Kamp, M. A., Alpdogan, S., Tevoufouet, E. E., Neiss, W. F., Hescheler, J., & Schneider, T. (2013). Ca 2.3 (R-type) calcium channels are critical for mediating anticonvulsive and neuroprotective properties of lamotrigine in vivo. Epilepsia. https://doi.org/10.1111/epi.12250

Eyo, U. B., Murugan, M., & Wu, L. J. (2017). Microglia-Neuron Communication in Epilepsy. Glia, 65(1), 5–18. https://doi.org/10.1002/glia.23006

Feng, H. J., & Faingold, C. L. (2017). Abnormalities of serotonergic neurotransmission in animal models of SUDEP. Epilepsy Behav, 71(Pt B), 174–180. https://doi.org/10.1016/j.yebeh.2015.06.008

Friedman, D., Honig, L. S., & Scarmeas, N. (2012). Seizures and epilepsy in Alzheimer’s disease. CNS Neurosci Ther, 18(4), 285–294. https://doi.org/10.1111/j.1755-5949.2011.00251.x

Fung, S., Smith, C. L., Prater, K. E., Case, A., Green, K., Osnis, L., Winston, C., Kinoshita, Y., Sopher, B., Morrison, R. S., Garden, G. A., & Jayadev, S. (2020). Early-Onset Familial Alzheimer Disease Variant PSEN2 N141I Heterozygosity is Associated with Altered Microglia Phenotype. J Alzheimers Dis, 77(2), 675–688. https://doi.org/10.3233/JAD-200492

Galla, L., Redolfi, N., Pozzan, T., Pizzo, P., & Greotti, E. (2020). Intracellular Calcium Dysregulation by the Alzheimer’s Disease-Linked Protein Presenilin 2. Int J Mol Sci, 21(3). https://doi.org/10.3390/ijms21030770

Ge, M., Zhang, J., Chen, S., Huang, Y., Chen, W., He, L., & Zhang, Y. (2022). Role of Calcium Homeostasis in Alzheimer’s Disease. Neuropsychiatr Dis Treat, 18, 487–498. https://doi.org/10.2147/NDT.S350939

Gonzalez-Reyes, R. E., Nava-Mesa, M. O., Vargas-Sanchez, K., Ariza-Salamanca, D., & Mora-Munoz, L. (2017). Involvement of Astrocytes in Alzheimer’s Disease from a Neuroinflammatory and Oxidative Stress Perspective. Front Mol Neurosci, 10, 427. https://doi.org/10.3389/fnmol.2017.00427

Jayadev, S., Case, A., Alajajian, B., Eastman, A. J., Moller, T., & Garden, G. A. (2013). Presenilin 2 influences miR146 level and activity in microglia. J Neurochem, 127(5), 592–599. https://doi.org/10.1111/jnc.12400

Jayadev, S., Case, A., Eastman, A. J., Nguyen, H., Pollak, J., Wiley, J. C., Moller, T., Morrison, R. S., & Garden, G. A. (2010). Presenilin 2 is the predominant gamma-secretase in microglia and modulates cytokine release. PLoS One, 5(12), e15743. https://doi.org/10.1371/journal.pone.0015743

Jayadev, S., Leverenz, J. B., Steinbart, E., Stahl, J., Klunk, W., Yu, C. E., & Bird, T. D. (2010). Alzheimer’s disease phenotypes and genotypes associated with mutations in presenilin 2. Brain, 133(Pt 4), 1143–1154. https://doi.org/10.1093/brain/awq033

Knox KM, B. M., Smith, CL, Jayadev S, Barker-Haliski M. (2023). Chronic Seizures Induce Sex-Specific Cognitive Deficits with Loss of Presenilin 2 Function. Exp Neurol.

Knox, K. M., Zierath, D. K., White, H. S., & Barker-Haliski, M. (2021). Continuous seizure emergency evoked in mice with pharmacological, electrographic, and pathological features distinct from status epilepticus. Epilepsia. https://doi.org/10.1111/epi.17089

Kolodziejczak, M., Bechade, C., Gervasi, N., Irinopoulou, T., Banas, S. M., Cordier, C., Rebsam, A., Roumier, A., & Maroteaux, L. (2015). Serotonin Modulates Developmental Microglia via 5-HT2B Receptors: Potential Implication during Synaptic Refinement of Retinogeniculate Projections. ACS Chem Neurosci, 6(7), 1219–1230. https://doi.org/10.1021/cn5003489

Krabbe, G., Matyash, V., Pannasch, U., Mamer, L., Boddeke, H. W., & Kettenmann, H. (2012). Activation of serotonin receptors promotes microglial injury-induced motility but attenuates phagocytic activity. Brain Behav Immun, 26(3), 419–428. https://doi.org/10.1016/j.bbi.2011.12.002

Lehmann, L., Lo, A., Knox, K. M., & Barker-Haliski, M. (2021). Alzheimer’s Disease and Epilepsy: A Perspective on the Opportunities for Overlapping Therapeutic Innovation. Neurochem Res. https://doi.org/10.1007/s11064-021-03332-y

Li, R., & Buchanan, G. F. (2019). Scurrying to Understand Sudden Expected Death in Epilepsy: Insights From Animal Models. Epilepsy Curr, 19(6), 390–396. https://doi.org/10.1177/1535759719874787

Lin, T. W., Liu, Y. F., Shih, Y. H., Chen, S. J., Huang, T. Y., Chang, C. Y., Lien, C. H., Yu, L., Chen, S. H., & Kuo, Y. M. (2015). Neurodegeneration in Amygdala Precedes Hippocampus in the APPswe/ PS1dE9 Mouse Model of Alzheimer’s Disease. Curr Alzheimer Res, 12(10), 951–963. https://doi.org/10.2174/1567205012666151027124938

Loewen, J. L., Barker-Haliski, M. L., Dahle, E. J., White, H. S., & Wilcox, K. S. (2016). Neuronal Injury, Gliosis, and Glial Proliferation in Two Models of Temporal Lobe Epilepsy. J Neuropathol Exp Neurol, 75(4), 366–378. https://doi.org/10.1093/jnen/nlw008

Lopera, F., Marino, C., Chandrahas, A. S., O’Hare, M., Villalba-Moreno, N. D., Aguillon, D., Baena, A., Sanchez, J. S., Vila-Castelar, C., Ramirez Gomez, L., Chmielewska, N., Oliveira, G. M., Littau, J. L., Hartmann, K., Park, K., Krasemann, S., Glatzel, M., Schoemaker, D., Gonzalez-Buendia, L.,… Quiroz, Y. T. (2023). Resilience to autosomal dominant Alzheimer’s disease in a Reelin-COLBOS heterozygous man. Nat Med, 29(5), 1243–1252. https://doi.org/10.1038/s41591-023-02318-3

Massey, C. A., Thompson, S. J., Ostrom, R. W., Drabek, J., Sveinsson, O. A., Tomson, T., Haas, E. A., Mena, O. J., Goldman, A. M., & Noebels, J. L. (2021). X-linked serotonin 2C receptor is associated with a non-canonical pathway for sudden unexpected death in epilepsy. Brain Commun, 3(3), fcab149. https://doi.org/10.1093/braincomms/fcab149

Meeker, S., Beckman, M., Knox, K. M., Treuting, P. M., & Barker-Haliski, M. (2019). Repeated Intraperitoneal Administration of Low-Concentration Methylcellulose Leads to Systemic Histologic Lesions Without Loss of Preclinical Phenotype. J Pharmacol Exp Ther. https://doi.org/10.1124/jpet.119.257261

Mehta, D., Jackson, R., Paul, G., Shi, J., & Sabbagh, M. (2017). Why do trials for Alzheimer’s disease drugs keep failing? A discontinued drug perspective for 2010-2015. Expert Opin Investig Drugs, 26(6), 735–739. https://doi.org/10.1080/13543784.2017.1323868

Metaxas, A., Anzalone, M., Vaitheeswaran, R., Petersen, S., Landau, A. M., & Finsen, B. (2019). Neuroinflammation and amyloid-beta 40 are associated with reduced serotonin transporter (SERT) activity in a transgenic model of familial Alzheimer’s disease. Alzheimers Res Ther, 11(1), 38. https://doi.org/10.1186/s13195-019-0491-2

Minkeviciene, R., Ihalainen, J., Malm, T., Matilainen, O., Keksa-Goldsteine, V., Goldsteins, G., Iivonen, H., Leguit, N., Glennon, J., Koistinaho, J., Banerjee, P., & Tanila, H. (2008). Age-related decrease in stimulated glutamate release and vesicular glutamate transporters in APP/PS1 transgenic and wild-type mice. J Neurochem, 105(3), 584–594. https://doi.org/10.1111/j.1471-4159.2007.05147.x

Minkeviciene, R., Rheims, S., Dobszay, M. B., Zilberter, M., Hartikainen, J., Fulop, L., Penke, B., Zilberter, Y., Harkany, T., Pitkanen, A., & Tanila, H. (2009). Amyloid beta-induced neuronal hyperexcitability triggers progressive epilepsy. J Neurosci, 29(11), 3453–3462. https://doi.org/10.1523/JNEUROSCI.5215-08.2009

Petrucci, A. N., Joyal, K. G., Purnell, B. S., & Buchanan, G. F. (2020). Serotonin and sudden unexpected death in epilepsy. Exp Neurol, 325, 113145. https://doi.org/10.1016/j.expneurol.2019.113145

Pisani, A., Bonsi, P., Martella, G., De Persis, C., Costa, C., Pisani, F., Bernardi, G., & Calabresi, P. (2004). Intracellular calcium increase in epileptiform activity: modulation by levetiracetam and lamotrigine. Epilepsia, 45(7), 719–728. https://doi.org/10.1111/j.0013-9580.2004.02204.x

Ptak, K., Yamanishi, T., Aungst, J., Milescu, L. S., Zhang, R., Richerson, G. B., & Smith, J. C. (2009). Raphe neurons stimulate respiratory circuit activity by multiple mechanisms via endogenously released serotonin and substance P. J Neurosci, 29(12), 3720–3737. https://doi.org/10.1523/JNEUROSCI.5271-08.2009

Racine, R., Okujava, V., & Chipashvili, S. (1972). Modification of seizure activity by electrical stimulation. 3. Mechanisms. Electroencephalogr Clin Neurophysiol, 32(3), 295–299.

Revuelta, M., Urrutia, J., Villarroel, A., & Casis, O. (2022). Microglia-Mediated Inflammation and Neural Stem Cell Differentiation in Alzheimer’s Disease: Possible Therapeutic Role of KV1.3 Channel Blockade. Front Cell Neurosci, 16, 868842. https://doi.org/10.3389/fncel.2022.868842

Reyes-Marin, K. E., & Nunez, A. (2017). Seizure susceptibility in the APP/PS1 mouse model of Alzheimer’s disease and relationship with amyloid beta plaques. Brain Res, 1677, 93–100. https://doi.org/10.1016/j.brainres.2017.09.026

Richerson, G. B., & Buchanan, G. F. (2011). The serotonin axis: Shared mechanisms in seizures, depression, and SUDEP. Epilepsia, 52 *Suppl 1*, 28–38. https://doi.org/10.1111/j.1528-1167.2010.02908.x

Romero-Calvo, I., Ocon, B., Martinez-Moya, P., Suarez, M. D., Zarzuelo, A., Martinez-Augustin, O., & de Medina, F. S. (2010). Reversible Ponceau staining as a loading control alternative to actin in Western blots. Anal Biochem, 401(2), 318–320. https://doi.org/10.1016/j.ab.2010.02.036

Sano, F., Shigetomi, E., Shinozaki, Y., Tsuzukiyama, H., Saito, K., Mikoshiba, K., Horiuchi, H., Cheung, D. L., Nabekura, J., Sugita, K., Aihara, M., & Koizumi, S. (2021). Reactive astrocyte-driven epileptogenesis is induced by microglia initially activated following status epilepticus. JCI Insight, 6(9). https://doi.org/10.1172/jci.insight.135391

Tejani-Butt, S. M., Yang, J., & Pawlyk, A. C. (1995). Altered serotonin transporter sites in Alzheimer’s disease raphe and hippocampus. Neuroreport, 6(8), 1207–1210. https://doi.org/10.1097/00001756-199505300-00033

Tort-Merino, A., Falgas, N., Allen, I. E., Balasa, M., Olives, J., Contador, J., Castellvi, M., Junca-Parella, J., Guillen, N., Borrego-Ecija, S., Bosch, B., Fernandez-Villullas, G., Ramos-Campoy, O., Antonell, A., Rami, L., Sanchez-Valle, R., & Llado, A. (2022). Early-onset Alzheimer’s disease shows a distinct neuropsychological profile and more aggressive trajectories of cognitive decline than late-onset. Ann Clin Transl Neurol, 9(12), 1962–1973. https://doi.org/10.1002/acn3.51689

Tupal, S., & Faingold, C. L. (2021). Serotonin 5-HT(4) receptors play a critical role in the action of fenfluramine to block seizure-induced sudden death in a mouse model of SUDEP. Epilepsy Res, 177, 106777. https://doi.org/10.1016/j.eplepsyres.2021.106777

Turkin, A., Tuchina, O., & Klempin, F. (2021). Microglia Function on Precursor Cells in the Adult Hippocampus and Their Responsiveness to Serotonin Signaling. Front Cell Dev Biol, 9, 665739. https://doi.org/10.3389/fcell.2021.665739

van Dyck, C. H., Swanson, C. J., Aisen, P., Bateman, R. J., Chen, C., Gee, M., Kanekiyo, M., Li, D., Reyderman, L., Cohen, S., Froelich, L., Katayama, S., Sabbagh, M., Vellas, B., Watson, D., Dhadda, S., Irizarry, M., Kramer, L. D., & Iwatsubo, T. (2022). Lecanemab in Early Alzheimer’s Disease. New England Journal of Medicine, 388(1), 9–21. https://doi.org/10.1056/NEJMoa2212948

Vande Vyver, M., Barker-Haliski, M., Aourz, N., Nagels, G., Bjerke, M., Engelborghs, S., De Bundel, D., & Smolders, I. (2022a). Higher susceptibility to 6 Hz corneal kindling and lower responsiveness to antiseizure drugs in mouse models of Alzheimer’s disease. Epilepsia, 63(10), 2703–2715. https://doi.org/10.1111/epi.17355

Vande Vyver, M., Barker-Haliski, M., Aourz, N., Nagels, G., Bjerke, M., Engelborghs, S., De Bundel, D., & Smolders, I. (2022b). Higher susceptibility to 6 Hz corneal kindling and lower responsiveness to antiseizure drugs in mouse models of Alzheimer’s disease. Epilepsia. https://doi.org/10.1111/epi.17355

Vezzani, A. (2005). Inflammation and epilepsy. Epilepsy Curr, 5(1), 1–6. https://doi.org/10.1111/j.1535-7597.2005.05101.x

Vezzani, A., Balosso, S., & Ravizza, T. (2012). Inflammation and epilepsy. Handb Clin Neurol, 107, 163–175. https://doi.org/10.1016/B978-0-444-52898-8.00010-0

von Linstow, C. U., Waider, J., Bergh, M. S., Anzalone, M., Madsen, C., Nicolau, A. B., Wirenfeldt, M., Lesch, K. P., & Finsen, B. (2022). The Combined Effects of Amyloidosis and Serotonin Deficiency by Tryptophan Hydroxylase-2 Knockout Impacts Viability of the APP/PS1 Mouse Model of Alzheimer’s Disease. J Alzheimers Dis, 85(3), 1283–1300. https://doi.org/10.3233/JAD-210581

Vossel, K., Ranasinghe, K. G., Beagle, A. J., La, A., Ah Pook, K., Castro, M., Mizuiri, D., Honma, S. M., Venkateswaran, N., Koestler, M., Zhang, W., Mucke, L., Howell, M. J., Possin, K. L., Kramer, J. H., Boxer, A. L., Miller, B. L., Nagarajan, S. S., & Kirsch, H. E. (2021). Effect of Levetiracetam on Cognition in Patients With Alzheimer Disease With and Without Epileptiform Activity: A Randomized Clinical Trial. JAMA Neurol, 78(11), 1345–1354. https://doi.org/10.1001/jamaneurol.2021.3310

Yang, F., Chen, L., Yu, Y., Xu, T., Chen, L., Yang, W., Wu, Q., & Han, Y. (2022). Alzheimer’s disease and epilepsy: An increasingly recognized comorbidity. Front Aging Neurosci, 14, 940515. https://doi.org/10.3389/fnagi.2022.940515

Zawar, I. (2022). Seizures in Dementia Are Associated with Worse Clinical Outcomes, Higher Mortality Rates and Shorter Lifespans. American Epilepsy Society Annual Meeting, Nashville, TN, USA.

Zawar, I., & Kapur, J. (2023). Does Alzheimer’s disease with mesial temporal lobe epilepsy represent a distinct disease subtype? Alzheimers Dement. https://doi.org/10.1002/alz.12943

Zhang, Y., Kecskes, A., Copmans, D., Langlois, M., Crawford, A. D., Ceulemans, B., Lagae, L., de Witte, P. A., & Esguerra, C. V. (2015). Pharmacological characterization of an antisense knockdown zebrafish model of Dravet syndrome: inhibition of epileptic seizures by the serotonin agonist fenfluramine. PLoS One, 10(5), e0125898. https://doi.org/10.1371/journal.pone.0125898

Zheng, X. Y., Zhang, H. C., Lv, Y. D., Jin, F. Y., Wu, X. J., Zhu, J., & Ruan, Y. (2022). Levetiracetam alleviates cognitive decline in Alzheimer’s disease animal model by ameliorating the dysfunction of the neuronal network. Front Aging Neurosci, 14, 888784. https://doi.org/10.3389/fnagi.2022.888784

Ziyatdinova, S., Gurevicius, K., Kutchiashvili, N., Bolkvadze, T., Nissinen, J., Tanila, H., & Pitkanen, A. (2011). Spontaneous epileptiform discharges in a mouse model of Alzheimer’s disease are suppressed by antiepileptic drugs that block sodium channels. Epilepsy Res, 94(1-2), 75–85. https://doi.org/10.1016/j.eplepsyres.2011.01.003

Ziyatdinova, S., Viswanathan, J., Hiltunen, M., Tanila, H., & Pitkanen, A. (2015). Reduction of epileptiform activity by valproic acid in a mouse model of Alzheimer’s disease is not long-lasting after treatment discontinuation. Epilepsy Res, 112, 43–55. https://doi.org/10.1016/j.eplepsyres.2015.02.005

