## Supplemental for "Chronic evoked seizures in young pre-symptomatic APP/PS1 mice induce serotonin changes and accelerate onset of Alzheimer’s disease-related neuropathology"

### Supplementary material

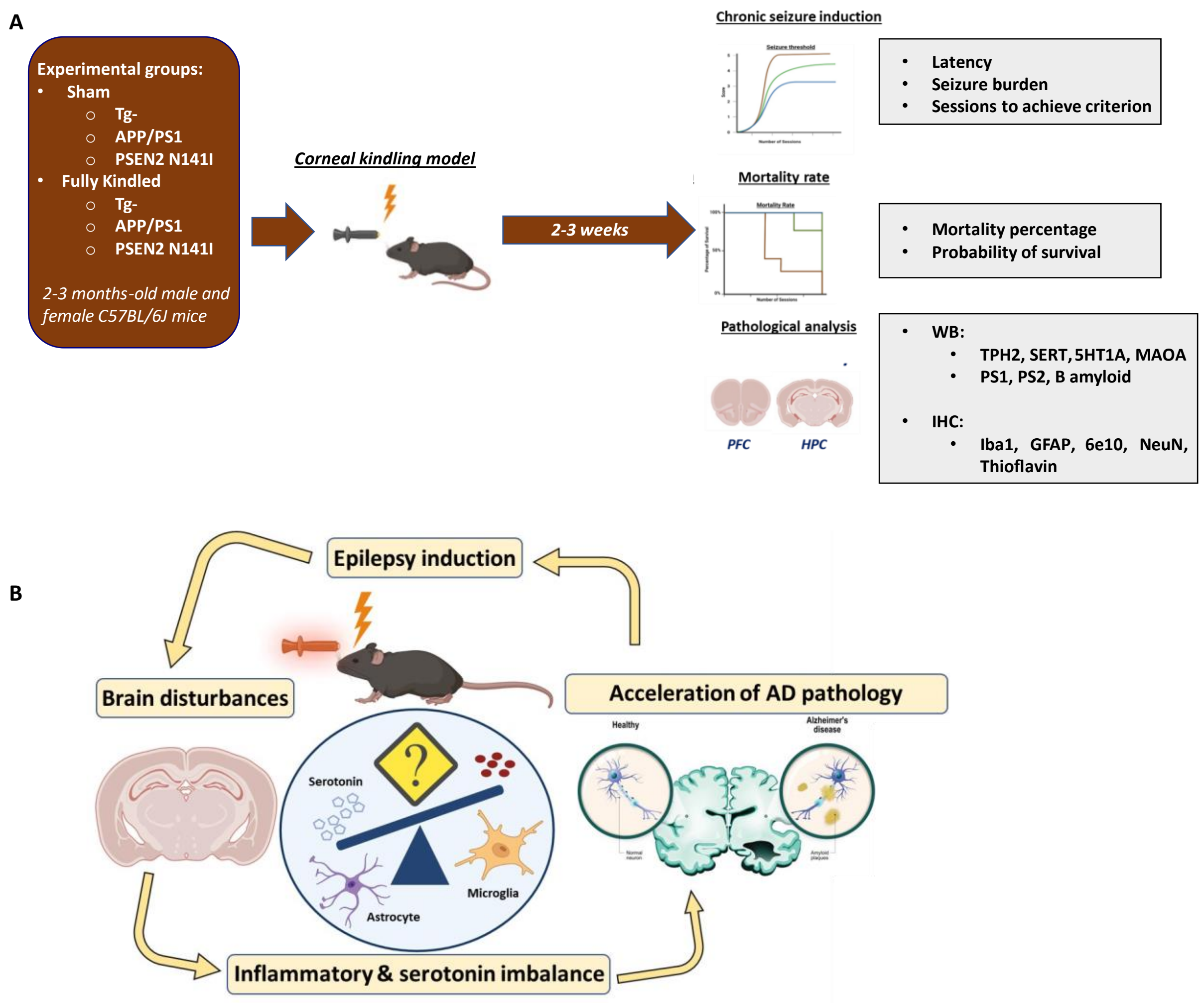


**Supplementary Figure S1. A)** Outline of the experimental design. **B)** Illustration of the potential interaction between kindling, neuroinflammation and gliosis, and the 5-HT pathway in AD pathology.

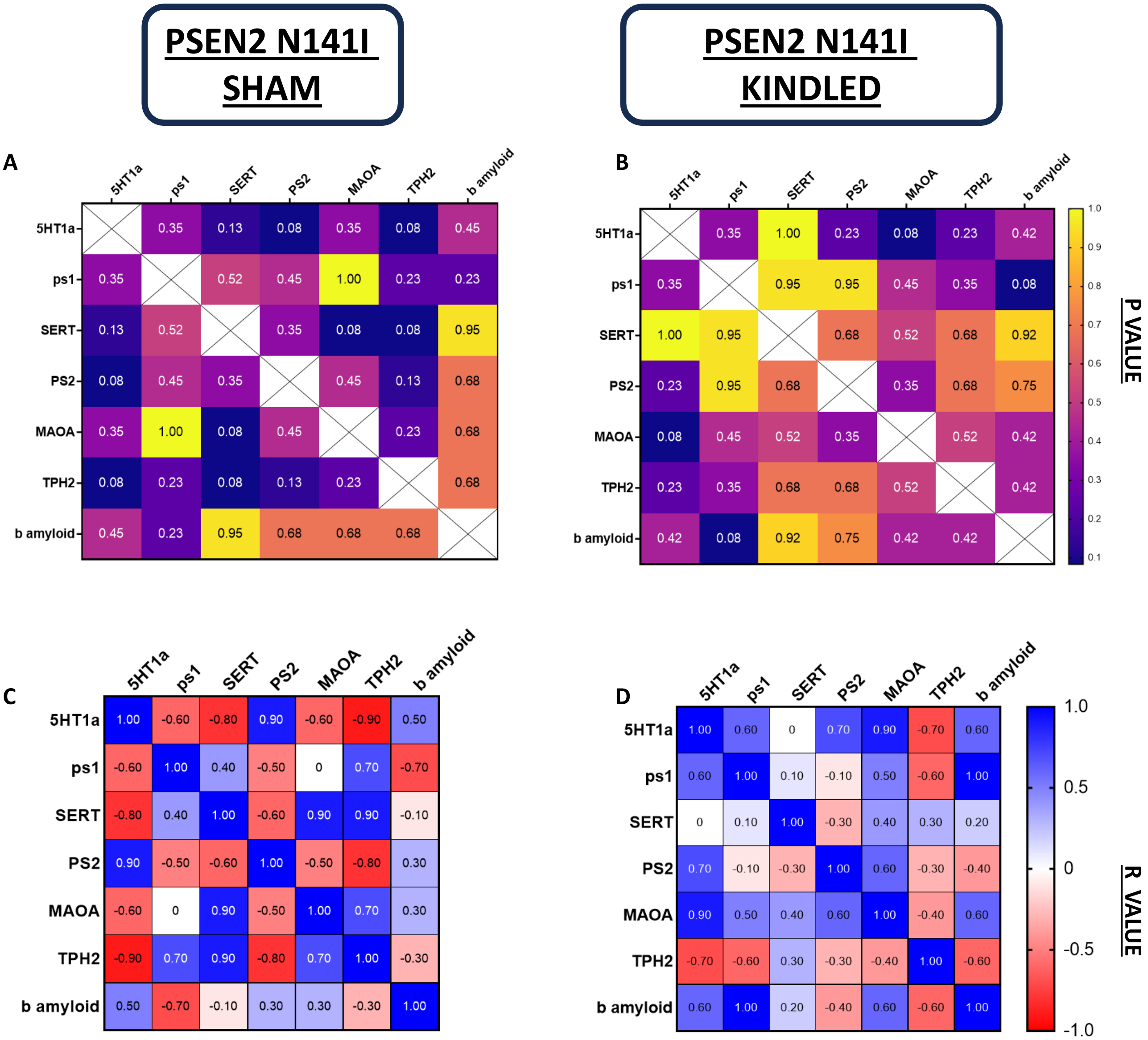


**Supplementary Figure S2.** Both PSEN2 N141I sham (**A, C**) and kindled (**B, D**) did not show any hippocampal correlations between 5-HT and AD-related proteins.

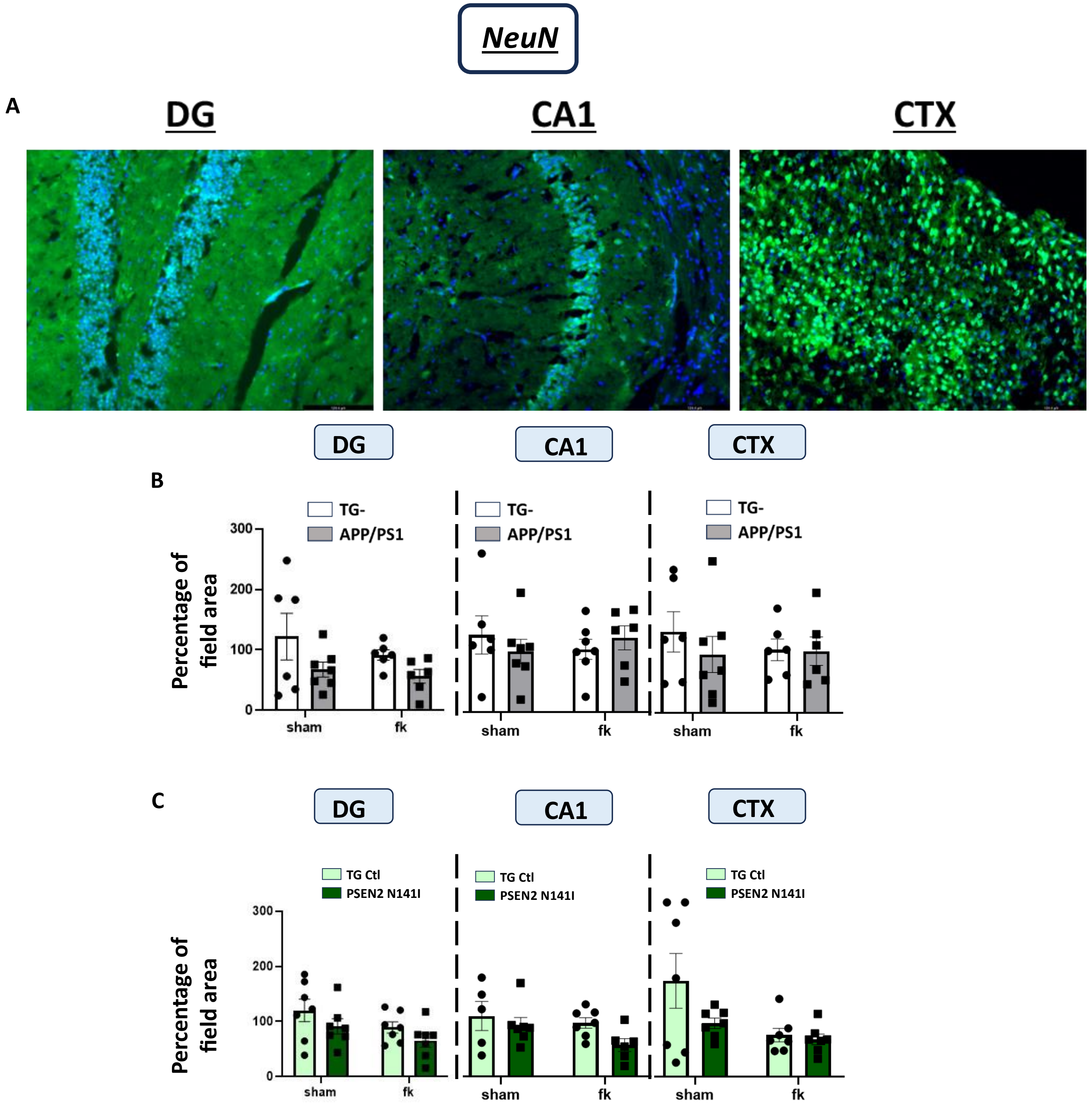


**Supplementary Figure s3. NeuN immunoreactivity in DG, CA1 and CTX. A)** Representative images of DAPI+NeuN staining. No significative differences were appreciated in either the **B)** APP/PS1 or **C)** PSEN2-N141I sham/kindled mice compared to the controls. Bars represent the median and the SEM. Data were assessed by two-way analysis of variance followed by the Bonferroni post-hoc test. N= 5-8.

| Group | Sex | Average weight (g) | Behavioral studies | | Molecular studies | | |
|---|---|---|---|---|---|---|---|
| | | | Total fully kindled mice (of total enrolled) | Mortality of kindled mice | Total sham mice | Total fully kindled mice (of total enrolled) | Mortality of kindled mice |
| Tg- | Male | 23.28 | 24/24 | 3/24 | 6 | 7/7 | 0/7 |
| | Female | 18.38 | 27/27 | 3/25 | 6 | 7/7 | 0/7 |
| APP/PS1 | Male | 23.15 | 19/19 | 12/19 | 6 | 6/6 | 0/6 |
| | Female | 19.28 * | 17/17 | 12/17 | 8 | 9/9 | 7/9 |
| Tg Ctl | Male | 24.44 | 39/48 | 5/49 | 10 | 3/8 | 0/8 |
| | Female | 18.80 | 39/39 | 0/40 | 8 | 7/7 | 0/7 |
| PSEN2-N141I | Male | 24.79 | 29/30 | 1/31 | 5 | 7/7 | 0/7 |
| | Female | 18.92 | 19/20 | 0/20 | 6 | 5/5 | 0/5 |

**Supplemental Table S1.** Information of the mice used for molecular studies. * $p < 0.05$ vs Tg- Females

| | Antibody | Concentration | Ref # | Company |
|---|---|---|---|---|
| Western blot | Tryptophan Hydroxylase 2 | 1:400 | ab126751 | Abcam, Cambridge, UK |
| | Serotonin Transporter | 1:250 | 19559-1-AP | Proteintech Genomics, San Diego, CA |
| | 5-HT1A | 1:300 | ab85615 | Abcam, Cambridge, UK |
| | Monoamine Oxidase A | 1:500 | ab184505 | Abcam, Cambridge, UK |
| | Presenilin 1 | 1:1000 | ab76083 | Abcam, Cambridge, UK |
| | Presenilin 2 | 1:1000 | ab51249 | Abcam, Cambridge, UK |
| | β-amyloid (clone 6E10; residues 1-16) | 1:500 | SIG-39320 | Biolegend, San Diego, CA |
| IHC | Iba-1 | 1:1000 | 019-19741 | FUJIFILM Wako Chemicals |
| | Cy3-conjugated GFAP | 1:1000 | C9205 | Sigma Aldrich, Saint Louis, MO |
| | NeuN | 1:300 | MAB377X | Millipore, Burlington, MA |
| | Alexa 488 conjugated b-amyloid (clone 6E10; residues 1-16) | 1:600 | 803013 | Biolegend, San Diego, CA |
| | Thioflavin S | 1% | T3516 | Sigma Aldrich, Saint Louis, MO |

**Supplemental Table S2.** Information about the antibodies used to perform molecular analysis.
